# The paracaspase MALT1 controls cholesterol homeostasis in glioblastoma stem-like cells through lysosome proteome shaping

**DOI:** 10.1101/2023.02.27.530259

**Authors:** Clément Maghe, Kilian Trillet, Gwennan André-Grégoire, Mathilde Kerhervé, Laura Merlet, Kathryn A Jacobs, Nicolas Bidère, Julie Gavard

## Abstract

Glioblastoma stem-like cells (GSCs) compose a tumor-initiating and -propagating population, remarkably vulnerable to any variation in the stability and integrity of the endolysosomal compartment. Previous work showed that the expression and activity of the paracaspase MALT1 control GSC viability via lysosomal abundance. However, the underlying mechanisms remain elusive. By combining RNAseq with proteome-wide label-free quantification, we now report that MALT1 repression in patient-derived GSCs alters the cholesterol homeostasis, which aberrantly accumulates in lysosomes. This failure in cholesterol supply culminates in cell death and autophagy defects, which can be partially reverted by providing exogenous membrane-permeable cholesterol to GSCs. From a molecular standpoint, targeted lysosome proteome analysis unraveled that NPC lysosomal cholesterol transporters were exhausted when MALT1 was held in check. Accordingly, we found that hindering NPC1 and NPC2 phenocopies MALT1 inhibition. This supports the notion that GSC fitness relies on lysosomal cholesterol homeostasis.

## INTRODUCTION

Glioblastoma (GB) is the most common and aggressive primary brain tumor in adults, as the survival median remains around 15 months after diagnosis. Despite standard-of-care treatments consisting of maximal surgical resection of the tumor bulk followed by concomitant radio- and chemo-therapies, relapses remain inexorable^1–5^. In addition to the specificity of the brain microenvironment, the aggressiveness and celerity of the expansion of GB can be ascribed to the high inter- and intra-tumor heterogeneity, complexifying the development of targeted therapies^6–9^. Pioneering studies characterized the tumor-initiating and propagating potential of a subpopulation of GB cells harboring stem properties, hereafter referred to as glioblastoma stem-like cells (GSCs)^10–12^. GSCs were notably defined their ability to self-renew, resist to conventional therapies, and invade the healthy brain^13–16^. In this context, the search for GSC vulnerabilities might offer new opportunities for therapeutic development. In keeping with this idea, several studies have demonstrated that the endo-lysosomal compartment is intrinsically tightly regulated in GSCs, in order to maintain their stemness capacities and viability^1, 17–21^. A low amount of endo-lysosomes might indeed sustain signaling receptors at the plasma membrane of GSCs, therefore perpetuating their constant activity, even in the harsh tumor environment^1, 2^. In this context, it was reported that the expression of the RNA binding protein quaking (QKI) supervised endo-lysosome abundance in experimental models, including patient-derived materials and genetically-modified mice^1, 2^. While a low level of lysosomes may ultimately favor the survival and maintenance of GSCs in the tumor microenvironment, destabilizing endo-lysosomal homeostasis has proven efficient in halting GSC growth and triggering GB decline *in vivo*^1, 17^.

The activity of lysosomes relies on the acidic pH of their lumen and an extensive number and diversity of hydrolases, but also on the active export of the as-generated products associated with the recycling activity of lysosomes^22–24^. Alterations in such processes lead to the development of a class of heterogeneous diseases, termed lysosomal storage disorders (LSDs), characterized by an aberrant accumulation of undigested, possibly toxic, materials^25, 26^. Likewise, the precise mechanistic description of these diseases had lengthened the list of cellular functions linked to lysosomes^27^. Accordingly, lysosomes have been attributed crucial roles in nutrient and lipid sensing, and therefore metabolism^28^. Several proteins acting as sensors, including NPC1, LYCHOS, and SLC38A9, were described to gauge cholesterol levels in lysosomes and further transmit the information to the cell^29–31^. In this context, the serine/threonine kinase mTOR plays a pivotal role. Its anchorage to the lysosomal surface and interaction with various metabolites and lipid sensors orchestrate the activation of the mTORC1 complex, therefore regulating anabolic (translation, lipogenesis, nucleotide biosynthesis) and catabolic (autophagy) processes^28, 30, 32, 33^. From a different angle, lysosomes unleash and spread extracellular lipids and cholesterol in intracellular membranes, via the degradation of endocytosed lipoproteins (LDL)^34–37^. In keeping with this idea, mutations in the lysosomal-cholesterol transporters NPC1 and NPC2 have been linked to the onset of the Niemann-Pick type C (NPC) disease. NPC characterized by an abnormal accumulation of cholesterol in the lumen and the limiting membrane of the lysosomes, impairing neuronal functionalities to culminate in mild-to-severe neurological defects in patients^38, 39^. Despite the central role of NPC proteins in coordinating cholesterol export from the lysosomes, scarce studies examined their putative roles in brain cancer.

Cholesterol is the building block for membrane organization and signaling platforms^40, 41^. As such, its homeostasis is strictly regulated through actionable checkpoints, including sensors and transcription factors. Among them, the sterol regulatory element-binding protein-2 (SREBP2) transcription factor plays a critical role in this process. Through the indirect sensing of cholesterol levels in the endoplasmic reticulum, SREBP2 triggers the expression of proteins and enzymes involved in *de novo* cholesterol synthesis (mevalonate pathway) and uptake (LDLR) to cover cellular cholesterol deficits^40, 42, 43^. Counterbalancing mechanisms concomitantly occur, such as the down-regulation of cholesterol exporters (ABCA family of transporters) via the inhibition of the LXR and RXR transcription factors^40, 44^. Combined, this warrants a finely tuned regulation of intracellular cholesterol levels.

The implication of such regulatory pathways in GB has mostly been investigated by means of *in silico* analysis (TCGA)^45^ and expression pattern^46^. However, manipulating cholesterol concentration was not thoroughly explored in these brain tumors, as compared to other cancer types^47^. For example, simvastatin, a cholesterol-reducing agent, dampened essential pro-oncogenic pathways and showed promising results in a clinical trial for breast cancer patients^48^. Moreover, the group of Paul S. Mischel demonstrated that the modulation of intracellular cholesterol levels could impact GB cells’ fitness^49^. Indeed, the induction of the expression of ABCA1 cholesterol exporter by the mean of LXR activation lowered GB cell’s intracellular cholesterol concentration and led to their death without affecting normal brain cell functions, suggesting a non-oncogene addiction of GB cells for cholesterol^49^. However, further description is needed to clarify the dependency of GSCs on cholesterol availability.

In this context, our present study linked cholesterol homeostasis to the paracaspase MALT1, which negatively regulates the endo-lysosomal compartment and further impacts GSC cell viability^1^. MALT1 is an arginine protease assembled into the CBM (CARMA-BCL10-MALT1) complex upon antigen receptor engagement in immune cells, as well as downstream of G-Protein Coupled Receptors and Receptor Tyrosine Kinases in non-immune cells^50–52^. Previous work demonstrated the constitutive activity of MALT1 in GSCs and ascribed a dominant function for this protein in regulating GSC fitness and viability^1^. In this context, MALT1 was shown to negatively correlate with the activity of QKI, ultimately down-regulating the endo-lysosomal compartment^1, 2^. Inhibiting MALT1 unleashed QKI activity, leading to a massive accumulation of dysfunctional and aberrant lysosomes, responsible in part for GSC cell death. Because of the endo-lysosomal remodeling, the autophagic flux as well as the mTORC1 pathway were obstructed by MALT1 inhibition, reinforcing the link between MALT1 and lysosome fitness^1^. However, the precise modifications operated by MALT1 on the endo-lysosomal compartment remain to be elucidated.

Here, using combined RNAseq and quantitative proteomic analysis, we unraveled an underestimated function of MALT1 in regulating the arsenal of cholesterol transporters in lysosomes. We showed that halting MALT1 led to SREBP2 transcription factor activation, as a consequence of cholesterol need. Accordingly, bioavailable cholesterol significantly rescued from cell death induced by MALT1 blockage, underlying a strong dependency of GSCs toward a fine-tuned cholesterol concentration. Mechanistically, we found that hindering MALT1 drastically remodeled the lysosomal proteome of GSCs, with a reduced amount of the NPC1 and NPC2 transporters, ultimately leading to cholesterol sequestration and accumulation in this compartment. As a proof of concept, inhibiting NPC1 and NPC2 decreased GSC cell viability. Disrupting intracellular cholesterol trafficking via the modification of the endo-lysosomal compartment might therefore represent an attractive opportunity for GSC eradication.

## Results

### The Inhibition of MALT1 Triggers the SREBP2 Transcriptional Program in Glioblastoma Stem-like Cells

To investigate the molecular basis for MALT1 control of the endo-lysosome fitness^1, 2^, a dual approach of RNAseq and proteome-wide label-free quantification (LFQ) analysis was conducted in patient-derived GSCs treated with the MALT1 inhibitor mepazine (MPZ)^53, 54^ (Figure 1A). The terms ’’cholesterol biosynthesis’’, ’’regulation of cholesterol biosynthesis by SREBP’’, and ’’metabolism of steroids’’ were among the top up-regulated pathways identified within the RNAseq dataset (Figure 1B, Gene Expression Omnibus <u>GSE139018</u>). This strong enrichment of mRNA encoding cholesterol-related genes was further visualized by gene set enrichment analysis (GSEA) of the reactome ’’cholesterol biosynthesis’’ and the “Horton Sterol Responsive Element Binding Factor targets’’ (Figure 1B). Accordingly, the proteomic analysis highlighted an over-representation of proteins related to a ’’lipid’’ node, in addition to “spindle”, “actin”, and “RNA” associated networks (Figure 1C, Tables S1-2). As *de novo* cholesterol synthesis occurs primarily through the mevalonate, Kandutsch-Russell and Bloch pathways^55^, we compared the level of transcripts and proteins involved in these metabolic arms (Figure 1D). Remarkably, most of the enzymes identified with this dual-omic approach were significantly increased upon MALT1 inhibition (Figure 1D). This effect was further validated at the RNA level for 8 out of 10 enzymes, upon MALT1 pharmacological inhibition with two compounds (namely mepazine and MLT748)^54, 56^ and RNA interference with two independent duplexes (Figure 1E). We also confirmed these results on the HSD17B7 target in two other patient-derived GSCs (GSC#4 and GSC#6) upon mepazine challenge (Figure 1F).

**Figure 1.**
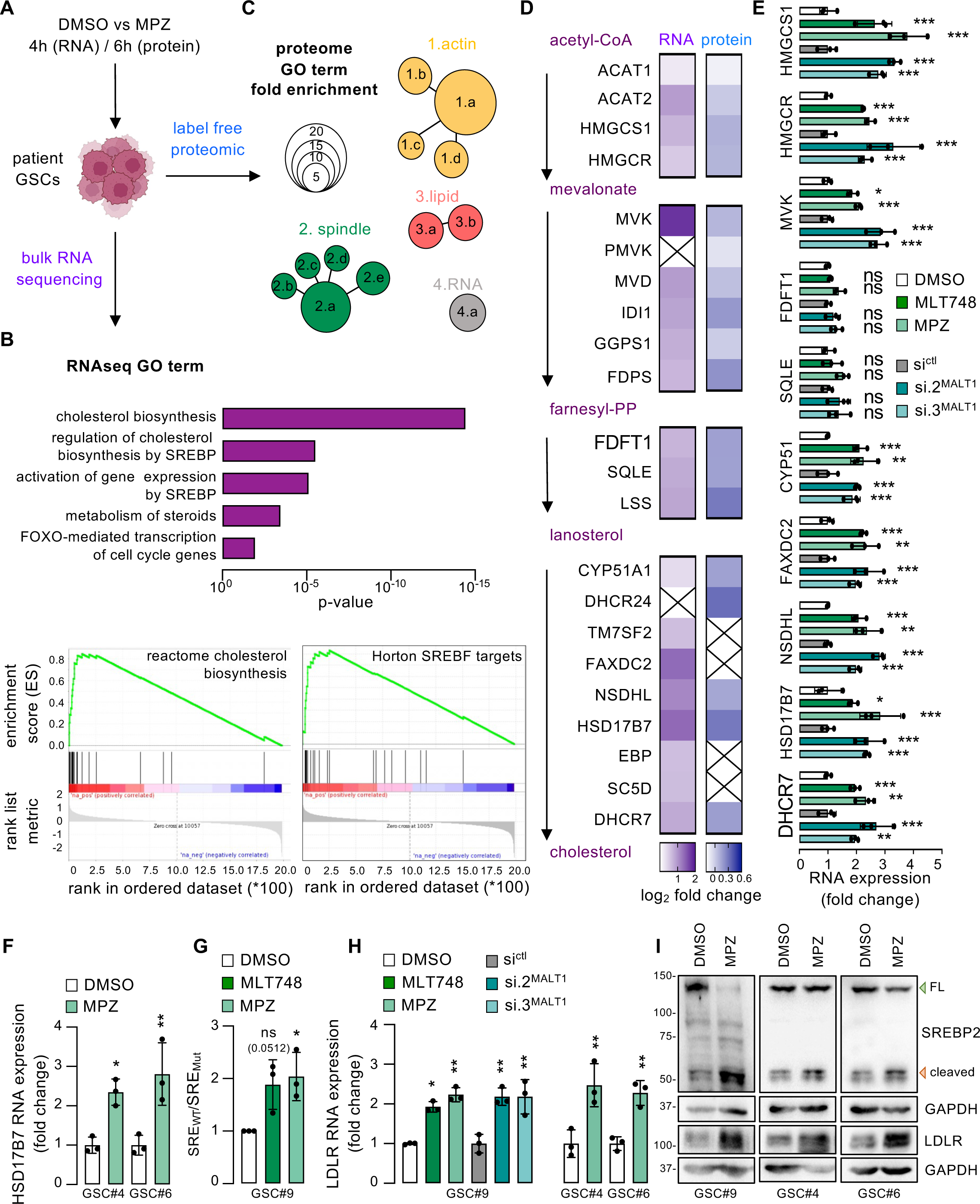
The Inhibition of MALT1 Triggers the SREBP2 Transcriptional Program in Glioblastoma Stem-like Cells. (A) Workflow of the dual approach to characterize the effects of MALT1 inhibition in patient-derived glioblastoma stem-like cells (GSCs). Patient-derived GSC#9 were treated with vehicle (DMSO) and the MALT1 inhibitor mepazine (MPZ, 20µM, 4h) and analyzed by RNA sequencing. Similarly, control and MPZ (20µM, 6h) treated GSC#9 were processed for proteome-wide label-free quantification (LFQ) analysis. (B) (Top) REAC enrichment analysis of the top upregulated pathways from RNAseq analysis of DMSO versus MPZ treated GSC#9. Upregulated genes (fold change > 1.5) upon MPZ treatment were analyzed. (Bottom) Gene set enrichment analysis (GSEA) plot of ‘’reactome cholesterol biosynthesis’’ and ‘’Horton SREBF targets’’ signatures. (C) Differentially upregulated proteins from DMSO versus MPZ-treated GSC#9 were analyzed with the Pantherdb^84^ pathway browser. Four main GO term signatures were identified as follows: 1. actin (1.a: actin filament network formation, 1.b: actin filament bundle assembly, 1.c: actin filament bundle organization, 1.d: positive regulation of actin filament polymerization), 2. spindle (2.a mitotic spindle organization, 2.b: mitotic spindle organization, 2.c: microtubule cytoskeleton organization involved in mitosis, 2.d: spindle assembly, 2.e: spindle organization), 3. lipid (3.a fatty acid oxidation, 3.b lipid oxidation), and, 4. RNA (4.a transcription elongation from RNA polymerase II promoter). (D) Heatmap of cholesterol synthesis gene expression (RNA, purple) and protein level (protein, blue) from GSC#9 treated as described in (A). The enzymes involved in the cholesterol synthesis pathway (black) with intermediate metabolites (purple) are shown on the left. RNA and protein selected hits from the RNAseq and proteome analysis are shown in log_2_ fold change. Non-identified genes and proteins are represented with a cross. Data are presented as the log_2_ fold change of MPZ vs DMSO treated GSC#9. RNAseq n=3, LFQ n=4. (E) qRT-PCR analysis of the indicated targets in GSC#9 treated for 4h with DMSO and MALT1 inhibitors (MPZ, 20µM and MLT748, 5µM). Alternatively, cells received non-silencing RNA duplexes (si^ctl^) and 2 independent duplexes targeting MALT1 (si.2^MALT1^ and si.3^MALT1^) for 3 days. Data were normalized to 2 housekeeping genes (HPRT1, ACTB) and are presented as the mean ± s.d. of 3 independent biological replicates. (F) qRT-PCR analysis of HSD17B7 in GSC#4 and GSC#6 treated as described in (E). Data were processed as described in (E). (G) GSC#9 were transfected with luciferase reporter plasmids for either wild-type SREBP2 promoter activity (SRE_WT_) or a mutated version (SRE_Mut_), to which SREBP2 cannot bind, in combination with Renilla under the control of neutral HSV-thymidine kinase promoter. GSC#9 were next treated for 16h with DMSO, MPZ (20µM), and MLT748 (5µM). Luminescence values were calculated as the ratio SRE_WT_/SRE_Mut_, and further normalized to Renilla intensities. Data are presented as the mean ± s.d. of 3 independent biological replicates. (H) qRT-PCR analysis of LDLR in GSC#9, GSC#4, and GSC#6, treated as described in (E). Data were processed as described in (E). (I) Western-blot analysis of the levels of SREBP2 and LDLR in GSC#9, GSC#4, and GSC#6, treated for 3h and 24h, respectively, with DMSO and MPZ (20µM). GAPDH served as a loading control. FL (full length) and cleaved SREBP2 forms are indicated with green and red arrowheads, respectively. All panels are representative of at least n=3, unless otherwise specified. t-test and ANOVA, *p<0.05, **p<0.01, ***p<0.001.

Most of the enzymes involved in *de novo* cholesterol biosynthesis are under the control of the SREBP2 transcription factor^57^ (Figure 1B). Interestingly, MALT1 inhibition with mepazine and MLT748 led to a two-fold increase in the SREBP2 promoter activity (Figure 1G). This was accompanied by an increase, both at the mRNA and the protein levels of LDLR, the main entry road for extracellular cholesterol, and a canonical SREBP2 target^34^, in three patient-derived cell lines (Figure 1H-I). Furthermore, *in silico* analysis of the TCGA database using the Gliovis^58^ tool uncovered the correlation between the *SREBP2* level of expression and the probability of survival in GB patients (Figure S1A). In fact, SREBP2 appeared significantly less expressed in GB samples. Of note, this was not the case for SREBP1 (Figure S1B), which operates in parallel with SREBP2 to regulate additional processes, such as lipid droplet generation and fatty acid biosynthesis^57^. Accordingly, single-cell RNAseq analysis^9^ also highlighted the accumulation of SREBP2-positive cells within the neoplastic cell and oligodendrocyte progenitor cell clusters, while SREBP1-positive cells were rather homogeneously distributed through cell entities, including myeloid cells (Figure S1C). SREBP2 is an endoplasmic reticulum resident protein that shuttles to the Golgi upon cholesterol depletion, where it is cleaved and subsequently enters the nucleus and acts as a transcriptional factor to cope with cholesterol deficits^42^. Western-blot analysis revealed robust processing of SREBP2 upon mepazine treatment in three patient-derived GSCs (Figure 1I and S1D). Further reinforcing this idea, the protein expression of the SREBP2 target LDLR was elevated when MALT1 was blocked using two separate pharmacological agents and RNA silencing (Figure 1I and S1D). By contrast, stopping MALT1 had no overt effect on the levels of DGAT1 (Figure S1E), an essential enzyme in triglyceride generation and lipid droplet assembling^59^, ruling out any possible bystander effect of MALT1 on non-cholesterol lipid homeostasis. Taken together, these results indicate that hampering MALT1 provokes the activation of the SREBP2 transcriptional program in GSCs, culminating in the transcription/translation of the enzymes involved in the synthesis and uptake of cholesterol.

### Cholesterol Abundance is Increased in MALT1-repressed GSCs

We next explored whether cell death resulting from MALT1 blockage^1^ could operate via SREBP2 activation. As expected, SREBP2 silencing precluded mepazine-associated increase in LDLR abundance (Figure S2A). However, cell death driven by mepazine was further augmented upon SREBP2 silencing (Figure S2B). Similar results were obtained with cerivastatin-induced inhibition of HMGCR, the rate-limiting enzyme in the cholesterol biosynthetic pathway^60^ (Figure S2C). Autophagy obstruction, as illustrated with the accumulation of P62 and LC3B lipidated forms, was also exacerbated when both MALT1 and SREBP2 were hampered (Figure S2D-E). Conversely, SREBP2 blockade alone was not sufficient to drive cell death and autophagy defects in GSCs (Figure S2B-E), suggesting rather an adaptive feedback mechanism for SREBP2 involvement in MALT1-related phenotypes than a direct downstream signaling cascade. The SREBP2 transcription factor function relies on small variations of cholesterol concentration in the endoplasmic reticulum^43^. Both the activation of SREBP2 and the aggravating action of SREBP2 silencing, therefore, suggest that GSCs face defective cholesterol level when MALT1 is held in check.

We next investigated whether the phenotypes caused by hampering MALT1 could reflect modifications in the cholesterol concentration and/or its handling in GSCs. Intracellular concentration in cholesterol is finely controlled by the *de novo* cholesterol synthesis, as well as uptake, storage, and export (Figure 2A). We first observed that MALT1 silencing resulted in elevated cholesterol concentration in total cell lysates (Figure 2B). Likewise, intracellular cholesterol content was further detected using the fluorescent polyene antibiotic filipin III that binds unesterified cholesterol^61^. Both flow cytometry and confocal analysis revealed a significant increase in filipin III staining (Figure 2C-D), therefore indicative of an overall raise in cholesterol content in response to MALT1 inhibition and silencing. LDL uptake through LDLR is one mechanism for transferring cholesterol. We, therefore, tracked fluorescently labeled LDL (dil-LDL) and found that MALT1-silenced GSCs significantly accumulated more LDL within two hours, possibly caused by the enhanced level of LDLR (Figures 1H-I, 2E). Same was true with the two MALT1 inhibitors. In contrast, the blockade of MALT1 in GSCs strongly reduced the expression of ABCA1, one of the main cholesterol exporters^40^ (Figure S2F). Hence, suppressing MALT1 caused GSCs to deploy an arsenal of strategies to increase total cholesterol concentration via synthesis and uptake while reducing its export.

**Figure 2.**
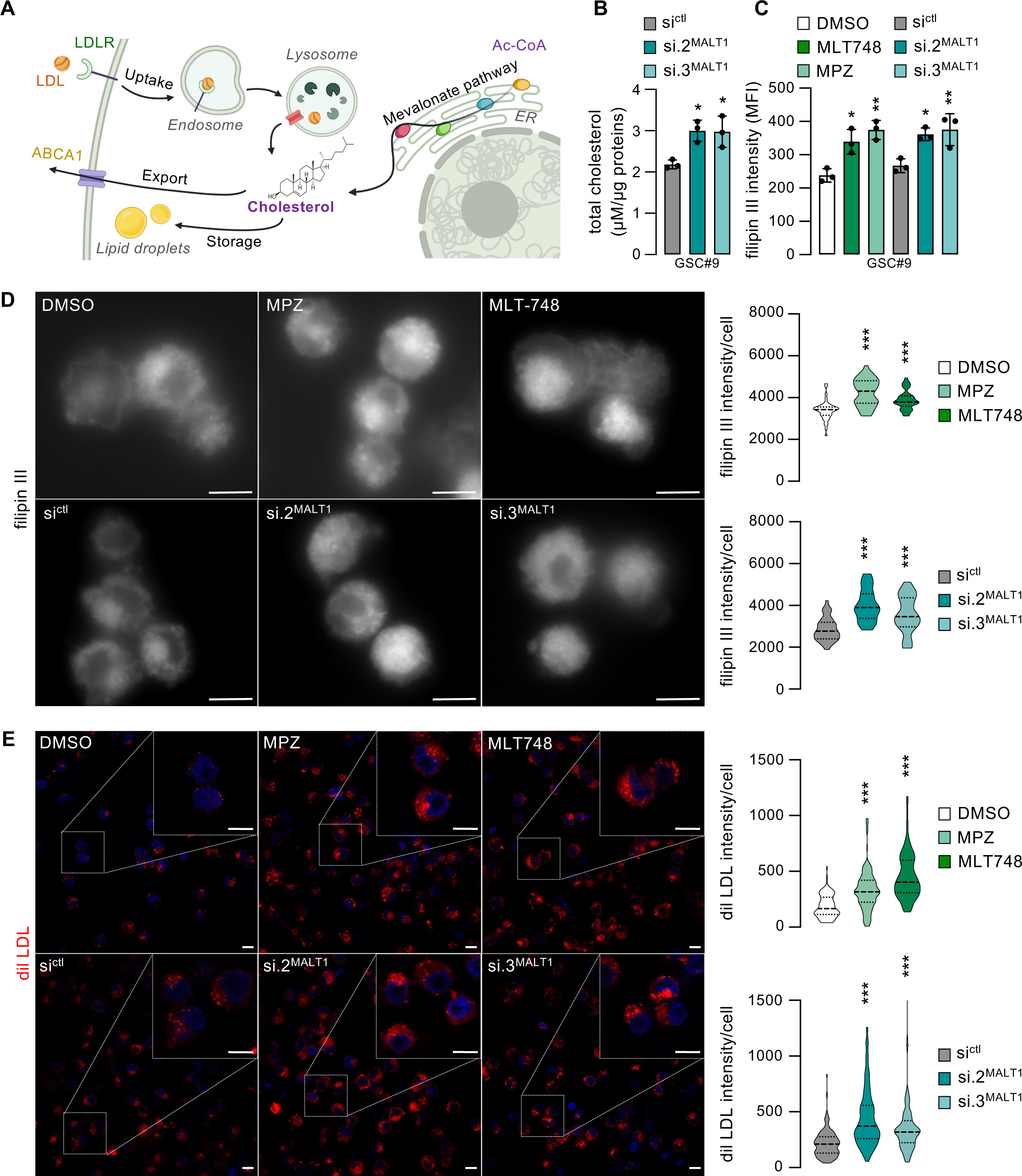
Cholesterol Abundance is Increased in MALT1-repressed GSCs. (A) Schematic representation of the different pathways which control the intracellular level of cholesterol via i) capture of cholesterol-filled LDL upon binding to the LDLR and trafficking through the endocytic/endosomal pathway, ii) *de novo* cholesterol synthesis into the mevalonate pathway, iii) handling the pool of free cholesterol via storage into lipid droplets, and, iv) export through different transporters such as ABCA1. (B) Total cholesterol level in GSC#9 transfected with non-silencing RNA duplexes (si^ctl^) and 2 independent duplexes targeting MALT1 (si.2^MALT1^ and si.3^MALT1^) for 3 days. Data are expressed as the ratio cholesterol/proteins (µM/µg) and presented as the mean ± s.d. of 3 independent biological replicates. (C) Flow cytometry analysis of the filipin III cholesterol probe in GSC#9 treated with vehicle (DMSO) and MALT1 Inhibitors (mepazine, MPZ, 20µM, and MLT748, 5µM) for 24h. Alternatively, cells received si^ctl^, si.2^MALT1^, and si.3^MALT1^ for 3 days. Data are presented as the mean ± s.d. of 3 independent biological replicates. (D) (Left) Confocal analysis of the filipin III cholesterol probe (gray) in GSC#9 treated as in (C). Scale bar: 10µm. (Right) Violin representation of the quantification of filipin III signal intensity per cell. n=38. (E) (Left) Confocal analysis of dil-LDL uptake (5µg/mL, 2h, red) in GSC#9 treated as in (C) for 16h. Nuclei are shown in blue (DAPI). Scale bar: 10µm. (Right) Violin representation of the quantification of dil-LDL signal intensity per cell. n>70. All panels are representative of at least n=3, unless otherwise specified. t-test and ANOVA, *p<0.05, **p<0.01, ***p<0.001.

### Bioavailable Cholesterol partially Counteracts MALT1 Inhibition-induced Cell Death

Because SREBP2 silencing further aggravated MALT1-based cell death (Figure S2B), we inferred that GSCs encountered a defective distribution of intracellular cholesterol, in spite of its apparent global accumulation. To challenge this hypothesis, cell viability was estimated in mepazine-treated GSCs, upon cholesterol feeding with free cholesterol or membrane-permeable cholesterol, *i.e.* coupled to MβCD . As expected, the depletion of cellular cholesterol with MβCD killed patient-derived GSCs in culture (Figure 3A). Mepazine-driven GSC demise could be significantly rescued using cholesterol/MβCD complexes, but not free cholesterol in three patient-derived GSCs (Figure 3A). In opposition to this, neither free nor complexed cholesterol protected GSCs from cell death driven by two classical lysosomal-destabilizing drugs, namely LLOMe and clemastine^17^(Figure 3B). This was also true in cells exposed to the mitochondrial-mediated intrinsic apoptosis activator, raptinal^62^ (Figure 3B), attributing thereby a specific effect of cholesterol supplementation on MALT1 actions. As expected, the levels of SREBP2 cleavage and of its downstream target LDLR were restored back upon cholesterol/MβCD addition in MALT1-inhibited GSCs (Figures 3C and S3A). Altogether, these data demonstrated that bioavailable cholesterol specifically rescued MALT1-induced cell death.

**Figure 3.**
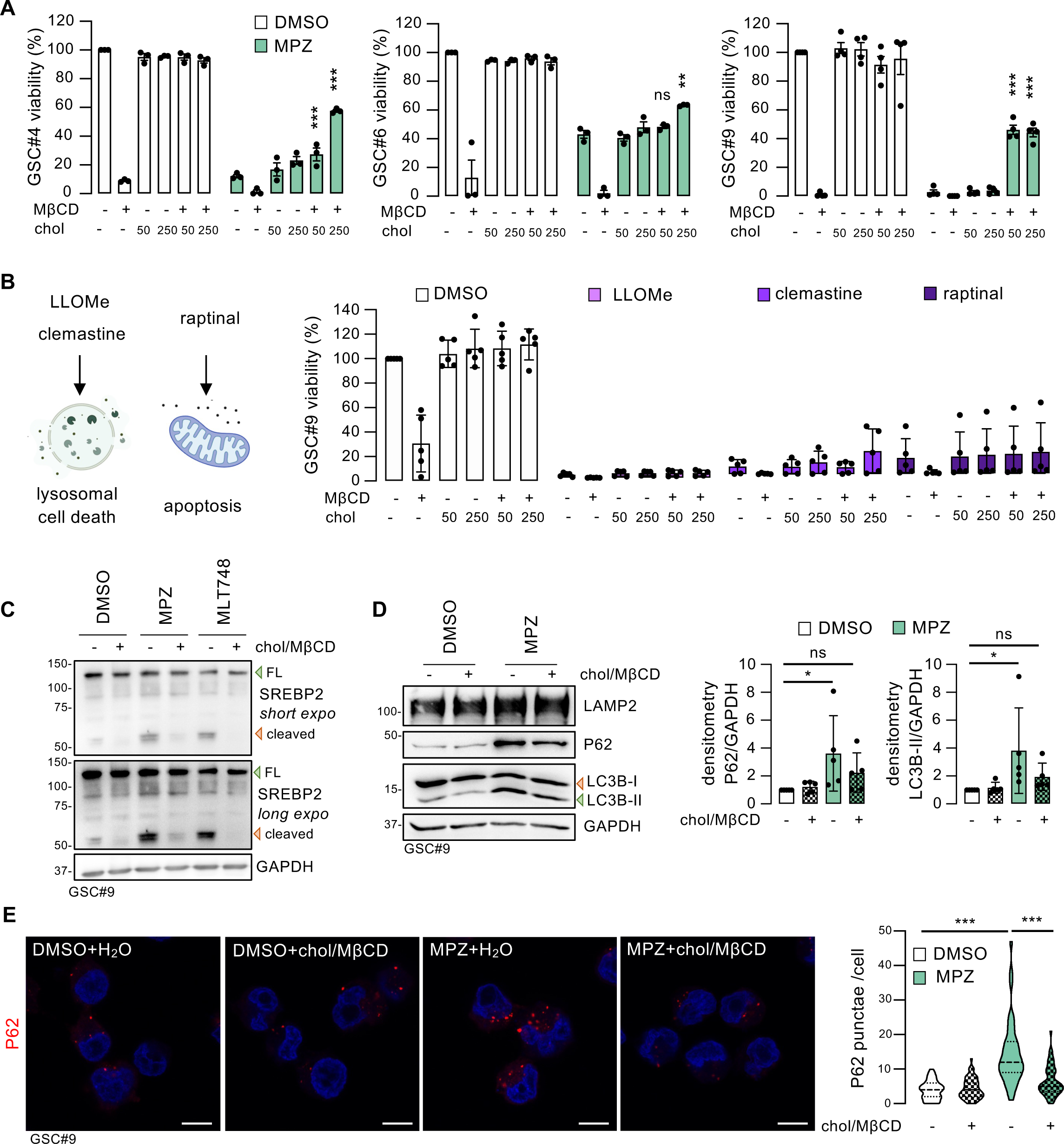
Bioavailable Cholesterol Partially Counteracts MALT1 Inhibition-induced Cell Death. (A) Cell viability was measured using CellTiter-Glo assay in GSC#4, GSC#6, and GSC#9 that were pre-treated for 1h with vehicle (DMSO) and mepazine (MPZ, 20µM), and then further challenged for 48h with vehicle (H_2_O), MβCD (0.1%), and either cholesterol alone (50 and 250µM) or in complex with Mβ . Data are presented as the mean ± s.d. of at least 3 independent biological replicates. Stars refer to the comparison with the MPZ pretreated and vehicle-challenged condition. (B) (Left) Schematic representation of the cell death processes induced by two lysosomal-destabilizing drugs (LLOMe and clemastine), and by the mitochondrial-mediated intrinsic apoptosis activator (raptinal). (Right) Cell viability was measured using CellTiter-Glo assay in GSC#9 that were pre-treated for 1h with DMSO, LLOMe (1µM), clemastine (20µM), and raptinal (2µM), and then further challenged for 48h with H_2_O, Mβ D (0.1%), and either cholesterol alone (50 and 250µM) or in complex with MβCD. Data are presented as the mean ± s.d. of 5 independent biological replicates. (C) Western-blot analysis of the level of SREBP2 in GSC#9 that were pre-treated for 1h with DMSO and MALT1 inhibitors (MPZ,20µM, and MLT748, 5µM), and then further challenged for 3h with vehicle (H_2_O) and with MβCD-complexed cholesterol (chol/MβCD, 250µM). GAPDH served as a loading control. FL (full length) and cleaved SREBP2 forms are indicated with green and red arrowheads, respectively. (D) (Left) Western-blot analysis of the levels of LAMP2, P62, and LC3B in GSC#9 treated as described in (C) for 24h. GAPDH served as a loading control. Unlipidated (LC3B-I) and lipidated (LC3B-II) LC3B forms are indicated with red and green arrowheads, respectively. (Right) Densitometric analysis of the level of P62 and LC3B-II (lipidated form) normalized to GAPDH. Data are presented as the mean ± s.d. of 5 independent experiments. (E) (Left) Confocal analysis of P62 staining (red) in GSC#9 treated as described in (D). Nuclei are shown in blue (DAPI). Scale bar: 10µm. (Right) Violin representation of the quantification of P62 punctae per cell. n>39. All panels are representative of at least n=3, unless otherwise specified. t-test and ANOVA, *p<0.05, **p<0.01, ***p<0.001.

We then explored at which step in the dying process, cholesterol supplementation rescued GSCs. First, even in the presence of exogenous cholesterol, MALT1 proteolytic activity remained blocked upon mepazine and MLT748 treatment, as visualized by the cleavage of its substrate HOIL1^50^ (Figure S3B), suggesting that cholesterol did not directly alter neither MALT1 activity nor its pharmacological inhibition. Next, while cholesterol/Mβ D addition failed to halt the endo-lysosomal increase observed upon MALT1 blockade (Figures 3D and S3C), the autophagic defects were toned down. Indeed, P62 accumulation and to a lesser extent LC3B lipidation, were reduced upon cholesterol/MβCD addition in the MALT1 inhibition context (Figure 3D-E). As the abundance of LAMP2 lysosomal protein was not normalized upon cholesterol addition, the supplementation may act downstream of the lysosomal changes observed upon MALT1 suppression. Nonetheless, the autophagy-associated marks and cell viability were recovered, suggesting that the addition of exogenous cholesterol might circumvent the lysosome-based defects.

### MALT1 Inhibition Edits the Lysosomal Proteome of GSCs and Affects the Lysosomal Cholesterol Export Machinery

Lysosomes are important hubs for cholesterol sensing, processing, and delivery to other cellular compartments^30, 35, 36, 63^. Because cholesterol addition was able to rescue several phenotypes resulting from MALT1 blockade, this raised the question of the ability of lysosomes to correctly convey cholesterol when MALT1 was repressed. To gain further insight into the lysosomal modifications engendered by MALT1 inhibition, lysosomes were immunopurified, a technique called LysoIP^64^ (Figure 4A). To this end, a patient-derived cell line was engineered to stably express a HA- and a FLAG-tagged lysosomal protein TMEM192 (hereinafter referred to as HA-lyso and FLAG-lyso, respectively)(Figure 4B-C). LysoIP allowed the expected enrichment of the intact endo-lysosomal compartment in GSCs, as observed with the lysosomal membrane protein LAMP2 and the intraluminal protein cathepsin D (CTSD), in the absence of proteins typically resident of other organelles, namely EEA1 for endosomes, calreticulin for ER, GM130 for Golgi, and VDAC for mitochondria (Figure 4D).

**Figure 4.**
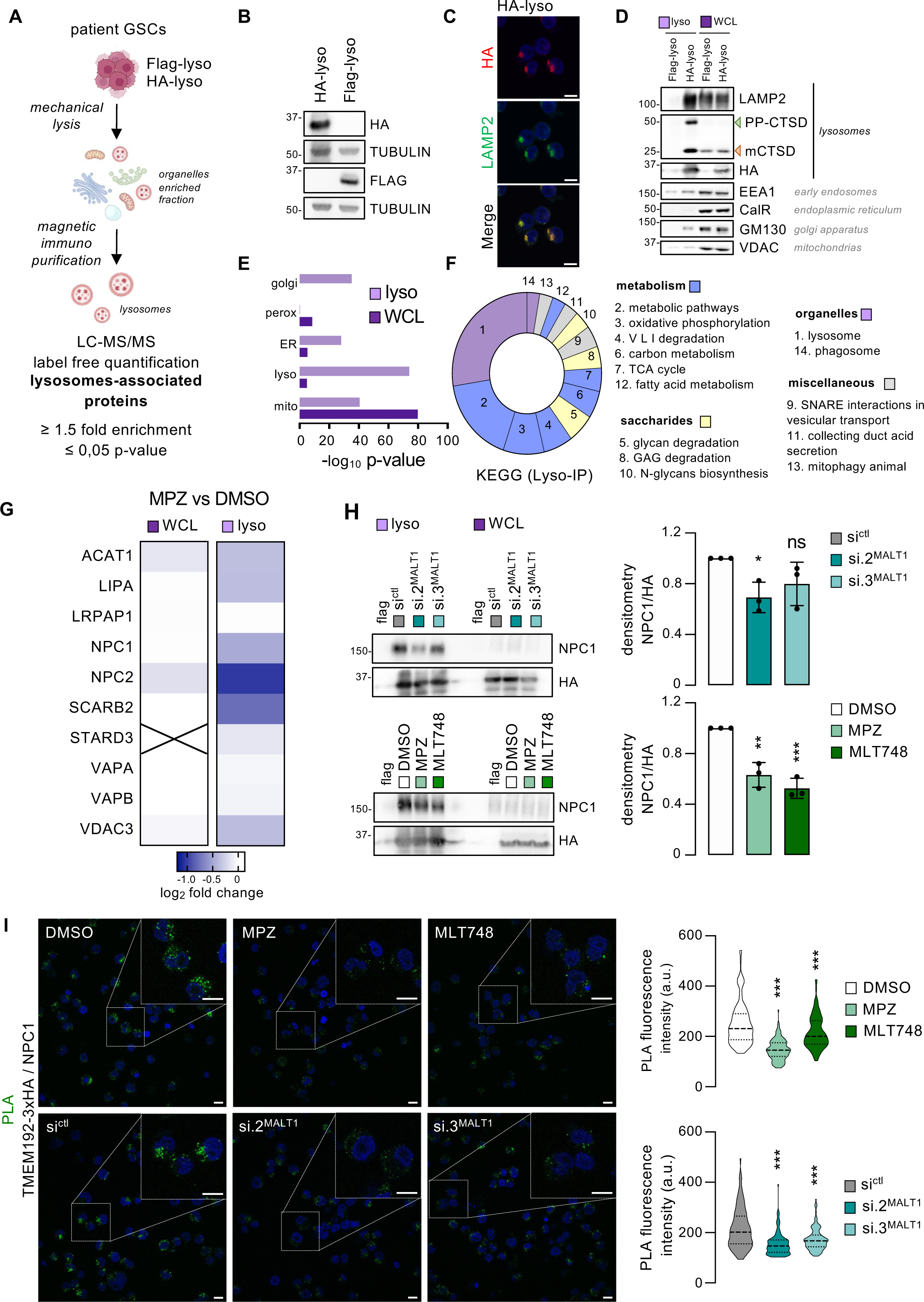
MALT1 Edits the Lysosomal Proteome of GSCs and Affects the Lysosomal Cholesterol Export Machinery. (A) Workflow of the Lyso-IP approach to characterize the effects of MALT1 inhibition on the lysosomal proteome in patient-derived GSCs. GSC#9 stably expressing the lysosomal protein TMEM192 tagged with either 3xHA (HA-lyso) or 2xFLAG (FLAG-lyso) were mechanically lysed. Lysosomes were immunoprecipitated with anti-HA magnetic beads. These immunopurified lysosome fractions were subsequently analyzed by western-blot and label-free quantification (LFQ) proteomic. Lysosome-associated proteins were defined as the hits with ≥1.5 fold change (HA/Flag) and p-value ≤ 05. (B) Western-blot analysis of the levels of TMEM192-3xHA (HA) and TMEM192-2xFLAG (FLAG) in lentivirally infected GSC#9. Tubulin served as a loading control. (C) Confocal analysis of TMEM192-3xHA (HA) staining (red) and LAMP2 staining (green) in HA-lyso GSC#9. Nuclei are shown in blue (DAPI). Scale bar: 10µm. (D) Western-blot analysis of the levels of LAMP2, CTSD (PP: pre-pro, m: mature), TMEM192-3xHA (HA), EEA1, CALRETICULIN (CalR), GM130, and VDAC in flag-lyso and HA-lyso GSC#9 immunopurified lysosomes (lyso, light purple square) and whole cell lysate (WCL, dark purple square). Organelles to which belong the proteins are indicated on the right. (E) Comparison of the GO:CC enrichment analysis of organelles (Golgi, peroxisomes, endoplasmic reticulum, lysosomes, and mitochondria) between WCL (dark purple square) and immunopurified lysosomes (lyso, light purple square). -log_10_ p-values are reported. (F) KEGG enrichment analysis of proteins defined as lysosomes-associated proteins (fold change >1.5 and p-value ≤0.05) in HA-lyso GSC#9 immunopurified lysosomes as compared to FLAG-lyso immunopurified lysosomes. (G) Heatmap of KEGG: Cholesterol Metabolism candidates from WCL (dark purple square) and lysosomes (lyso, light purple square) in HA-lyso GSC#9, treated with vehicle (DMSO) and mepazine (MPZ, 20µM, 6h). Data are presented as the log_2_ fold change. Candidates not identified are shown with a cross. (H) (Left) Western-blot analysis of the level of NPC1 from FLAG-lyso and HA-lyso GSC#9 immunopurified lysosomes (lyso, light purple square) and WCL (whole cell lysate, dark purple square). (Top) FLAG-lyso and HA-lyso GSC#9 were transfected with non-silencing RNA duplexes (si^ctl^) and 2 independent duplexes targeting MALT1 (si.2^MALT1^ and si.3^MALT1^) for 3 days. (Bottom) FLAG-lyso and HA-lyso GSC#9 were treated with DMSO, MPZ (20µM), and MLT748 (5µM) for 6h. TMEM192-3xHA (HA) served as a loading control. (Right) Densitometric analysis of NPC1 level normalized to TMEM192-3xHA (HA). Data are presented as the mean ± s.d. of 3 independent experiments. (I) (Left) Confocal analysis of Proximity Ligation Assay (PLA) between TMEM192-3xHA and NPC1. HA-lyso GSC#9 were treated as described in (H). Fluorescent signal (green) reflects a <40nM proximity. Nuclei are shown in blue (DAPI). Scale bar: 10µm. (Right) Violin representation of the quantification of the PLA signal intensity per cell. n>108. All panels are representative of at least n=3, unless otherwise specified. t-test and ANOVA, *p<0.05, **p<0.01, ***p<0.001.

The lysosomal proteome from vehicle- and mepazine-treated GSCs were then further inspected by label-free protein quantification (Figure 4A). The analysis established the purity degree and strong enrichment in lysosomes (Table S3). Indeed, unlike other organelles, the representation of lysosomal proteins was strongly increased in LysoIP samples, as compared to the whole cell lysate ones (WCL) (Figures 4E and S4A). Notably, a KEGG enrichment analysis showed a prominent signature of lysosomal proteins (Figure 4F). The comparison between the proteomes of lysosomes isolated from control and mepazine-treated GSCs highlighted autophagy defects (Figure S4C). As seen by a gene ontology:biological process (GO:BP) enrichment analysis, most of the proteins found enriched in lysosomes of MALT1-inhibited GSCs were related to autophagy (Figure S4B). The volcano plot visualization demonstrated a substantial accumulation of classical autophagic receptors, such as TAX1BP1 and SQSTM1 (P62) (Figure S4C). This initial examination of the lysosome proteome thus supports the idea that MALT1 inhibition is accompanied by a defect in the degradative capacity of lysosomes. Closer exploration into the differentially expressed proteins identified a sharp reduction in the level of most of the proteins related to cholesterol homeostasis, such as NPC1, NPC2, and SCARB2 transporters in the lysosomes, but not in whole cell lysate (Figure 4G). This is indicative of a probable change in the relative repartition of these lysosomal resident proteins. Accordingly, the mRNA level of NPC1 and NPC2 were left unchanged in response to MALT1 blockade (Figure S4E). This NPC1 deficit from lysosomes was then independently validated in cells challenged with MALT1 inhibitors and siRNA (Figure 4H). Moreover, proximity ligation assay (PLA) performed between NPC1 and TMEM192 revealed a loss in their association, thus suggesting that the number of NPC1-positive lysosomes was reduced when MALT1 was held in check (Figure 4I).

### NPC1 Blockade Partially Recapitulated MALT1-repressed Phenotypes in GSCs

We then tested whether NPC1 dilution from lysosomes could execute MALT1-related cell death in GSCs. *In silico* analysis of the TGCA database displayed that a low level of NPC1 RNA expression correlated with a significantly higher probability of survival in GB patients (Figure 5A). This was however not the case for NPC2 (Figure S5A). As expected, the use of U18666A, a classical NPC1 inhibitor^65^, promoted SREBP2 processing and LDLR expression^65^ (Figure 5B), paralleling the effects of MALT1 hindrance (Figure 1I). Likewise, the SREBP2 reporter assay supported the hypothesis of the activation of SREBP2 upon NPC1 blockade in GSCs (Figure 5C). Moreover, the filipin III staining was globally increased in NPC1-silenced GSCs (Figure 5D). This was further illustrated with filipin III-stained cholesterol that accumulated in lysosomes upon NPC1 inhibition and silencing (Figure 5E). Again, similar phenotype was observed in MALT1-arrested GSCs using drugs and RNA interference (Figure 5E). The level of the lipid lysobisphosphatidic acid (LBPA) that significantly accumulates in NPC1-inhibited cells (Figure 5F)^66, 67^, was also found to augment in response to MALT1 blockade, arguing for a shared response to MALT1 and NPC1 suppression. However, hindering NPC1 and NPC2 could not recapitulate the lysosomal increase observed when MALT1 activity/expression was abrogated (Figures 5H-I and S5B). The accumulation of the autophagic receptor P62 was however phenocopied (Figures 5H-I and S5B), suggesting that lysosomes from MALT1 and NPC1/2-inhibited GSCs might feature similar degradative defects. In this regard, U18666A treatment, as well as the silencing of NPC1 and NPC2 significantly reduced GSC viability (Figures 5J-K and S5C-D). Of interest, the reduction in cell viability upon U18666A treatment in non-GSC brain cells, such as human astrocytes and brain endothelial cells, was however not as toxic (Figure S5D). Likewise, lymphocytes without intrinsic MALT1 activity (*i.e.* Jurkat T cells and BJAB lymphoma) were left intact. Conversely, the viability of a MALT1-dependent OCI-Ly3 lymphoma cell line^68, 69^, was strongly affected by the U18666A treatment (Figure S5D), raising the possibility of a conserved role of MALT1 in cholesterol homeostasis.

**Figure 5.**
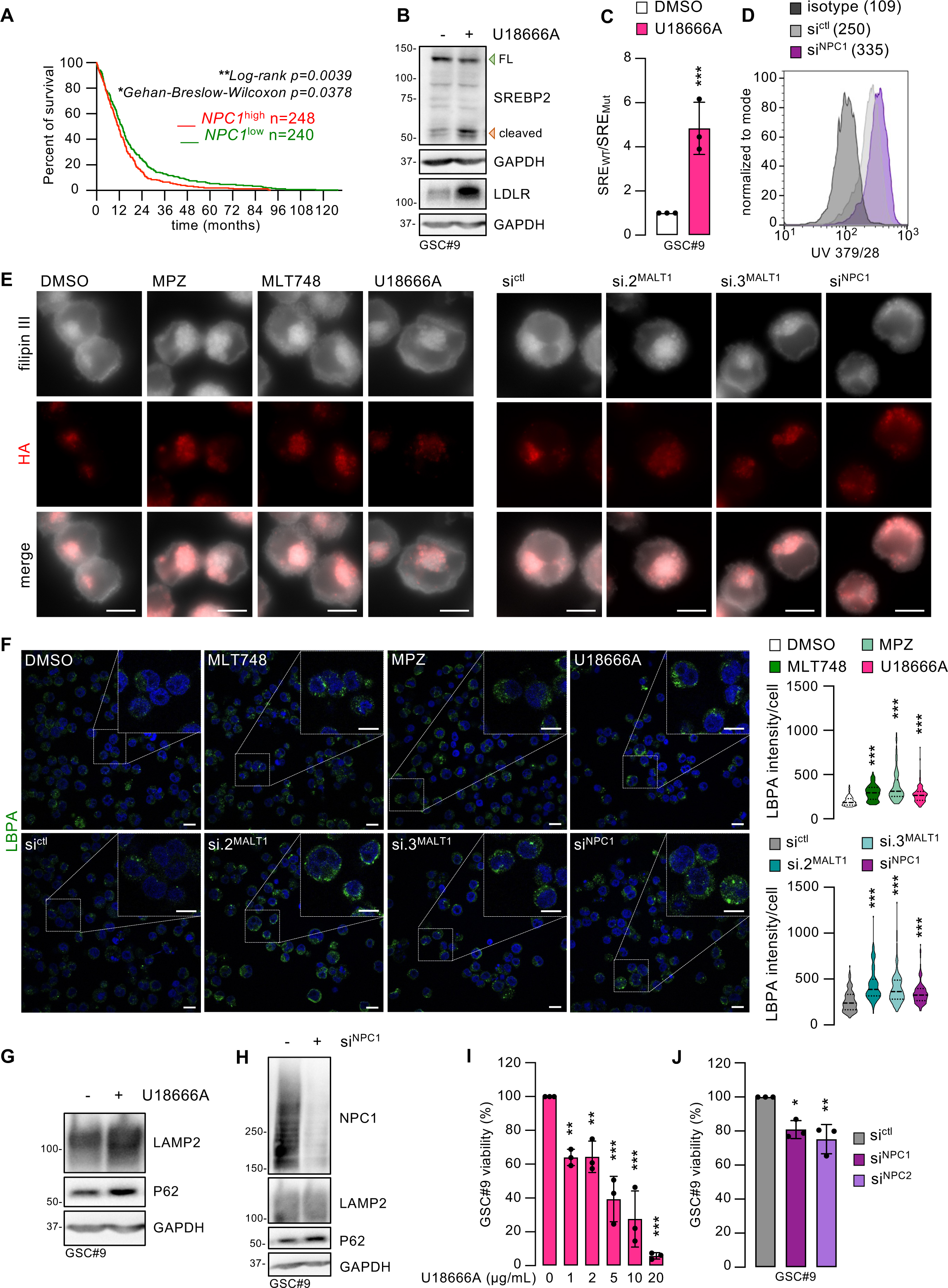
NPC1 Blockade Partially Recapitulated MALT1-repressed Phenotypes in GSCs. (A) Kaplan-Meier curve of the probability of survival for >240 GB patients with low (green) and high (red) *NPC1* RNA level based on the TCGA Agilent-4502A dataset. (B) Western-blot analysis of the levels of SREBP2 and LDLR in GSC#9 treated for 3h and 24h, respectively, with vehicle (DMSO) and U18666A (2µg/mL). GAPDH served as a loading control. FL (full length) and cleaved SREBP2 forms are indicated with green and red arrowheads, respectively. (C) GSC#9 were transfected with luciferase reporter plasmids for either wild-type SREBP2 promoter activity (SRE_WT_) or a mutated version (SRE_Mut_), in combination with TK-Renilla. GSC#9 were next treated for 16h with DMSO and U18666A (2µg/mL). Luminescence values were calculated as the ratio SRE_WT_/SRE_Mut_, and further normalized to Renilla intensities. Data are presented as the mean ± s.d. of 3 independent biological replicates. (D) Flow cytometry analysis of the filipin III cholesterol probe in GSC#9 transfected with non-silencing (si^ctl^) and NPC1 (si^NPC1^) targeting RNA duplexes for 3 days. Histogram is representative of at least 3 independent biological replicates. Mean fluorescence is indicated. E) Confocal analysis of cholesterol (filipin III) staining (gray) and TMEM192-3xHA (HA) staining (red) in HA-lyso GSC#9 treated with DMSO, MALT1 inhibitors (MPZ, 20µM and MLT748, 5µM), and NPC1 inhibitor (U18666A, 2µg/mL) for 16h. Alternatively, cells received si^ctl^, si^NPC1^, and 2 independent RNA duplexes targeting MALT1 (si.2^MALT1^ and si.3^MALT1^). Scale bar: 10µm. (F) (Left) Confocal analysis of lysobisphosphatidic acid (LBPA) staining (green) in GSC#9 treated for 24h as described in (E). Nuclei are shown in blue (DAPI). Scale bar: 10µm. (Right) Violin representation of the quantification of LBPA signal intensity per cell. n>112. (G) Western-blot analysis of the levels of LAMP2 and P62 in GSC#9 treated for 24h, as described in (C). GAPDH served as a loading control. (H) Western-blot analysis of the levels of NPC1, LAMP2, and P62 in GSC#9, transfected as in (D). GAPDH served as a loading control. (I) Cell viability was measured using CellTiter-Glo assay in GSC#9 treated for 48h with DMSO and U18666A at the indicated doses. Data are presented as the mean ± s.d. of 3 independent biological replicates. (J) Cell viability was measured using CellTiter-Glo assay in GSC#9 transfected for 3 days with si^ctl^, si^NPC1^, and si^NPC2^. Data are presented as the mean ± s.d. of 3 independent biological replicates. All panels are representative of at least n=3, unless otherwise specified. t-test and ANOVA, *p<0.05, **p<0.01, ***p<0.001.

## DISCUSSION

Taken together, our results support an underestimated role for the paracaspase MALT1 in the regulation of intracellular cholesterol homeostasis. Indeed, suppressing MALT1 activity/expression results in a profound remodeling of the lysosomal compartment accompanied by cholesterol retention in lysosomes and the subsequent failure in the intracellular delivery of this vital lipid. These defects ultimately led to the demise of glioblastoma stem-like cells.

First, hindering MALT1 provokes an accumulation in intracellular cholesterol, as estimated with direct cholesterol quantification and filipin III binding. Despite this apparent elevation in intracellular cholesterol levels, cells counterintuitively deploy a myriad of strategies usually in place to cope with cholesterol needs^40, 42^. This is notably shown by the RNAseq analysis of MALT1-blocked cells. Upon cholesterol depletion that it senses in the ER, SREBP2 boosts the expression of genes involved in cholesterol synthesis and uptake^43, 57^. Concomitantly, opposite effectors down-regulate cholesterol storage and export to maintain balanced intracellular cholesterol concentration^40, 44^. Our results, therefore, suggest that MALT1-inhibited GSCs execute a program to compensate for the retention of cholesterol in lysosomes. In a MALT1-negative context, hampering SREBP2-induced cholesterol synthesis with RNA interference or cerivastatin, an inhibitor of the rate-limiting enzyme of the cholesterol biosynthesis pathway^60^, aggravates both autophagy defects and cell death in GSCs. Thus, this suggests a strong dependency of these cells toward finely maintained cholesterol homeostasis. Abruptly lowering the cholesterol reserve appears as an efficient strategy for GSC elimination, as illustrated by the effect of the cholesterol-complexing agent MβD. Combining the targeting of these two non-oncogene addiction pathways, namely MALT1 and cholesterol level, may thus represent a valid strategy for GSC destruction. However, one caveat with the use of statins resides in the high number of reported negative actions on non-tumor cells, such as astrocytes^49, 70^. More specific cholesterol-lowering agents may be thus valuable. For example, LXR-623, a clinically viable and brain-penetrant LXR agonist can lower GB cell viability in a cholesterol-dependent fashion, without affecting healthy cells^49^.

A key point we address here concerns the mechanisms via which MALT1 blockage interrupts cholesterol supply. Our data identify that MALT1-off cells suffer from a lysosomal cholesterol-handling defect, most likely due to the reduced level of the main cholesterol transporters at the lysosomes. Notably, label-free quantitative proteomics unveils a reduction in NPC1 in these organelles while the overall expression remains steady in cells. Detailed biochemical and imaging studies of the lysosomal compartment upon MALT1 inhibition could help define the precise localization of NPC1. Paralleling the situation in Niemann-Pick type C-patients presenting mutations that might alter the folding of the protein or its ability to anchor in lipid-rich membranes, NPC1 could be retained in the ER^71, 72^. Although we cannot rule out that NPC1 is rerouted to different cellular membranes, it is conceivable that NPC1 is diluted in the pool of newly generated lysosomes. In keeping with this idea, MALT1 silencing might create NPC1-exhausted lysosomes, making the as-produced lysosome population less prone to export cholesterol while it continues to transit via these organelles^36, 37, 63, 73^. Arguing in favor of the apparition of a pool of aberrant and dysfunctional NPC1-defective lysosomes, targeted proteomics highlights an autophagy signature. This is in agreement with previous studies that demonstrate the robust autophagic defects in NPC1-null models^30, 36, 74, 75^. Moreover, as blocking SREBP2-induced cholesterol replenishment in MALT1-halted GSCs enhances autophagic failure, lysosomal-cholesterol handling, therefore, appears as a major actor in the autophagic pathway termination. However, how membrane permeable cholesterol addition onto MALT1-arrested GSCs rescues this phenotype is still unclear. Overall, the increased abundance of abnormal lysosomes results in a global accumulation and sequestration of cholesterol, subsequently involved in the induction of most of the MALT1 inhibition-induced phenotypes.

Although cholesterol is viewed as an essential metabolite fueling tumor cells^47–49^, how exactly the cholesterol inflation in lysosomes leads to cell death remains to be elucidated. Hints might come from the parallel with lysosomal storage disease, such as the Niemann-Pick type C disease. In NPC patient-derived cells and experimental models, cholesterol is trapped in lysosomes^30, 36, 38, 66, 67^, similar to our observations in MALT1-defective cells. It is noteworthy that MALT1 suppression reiterates NPC disease traits, including a massive accumulation of cholesterol in lysosomes, the fragility of the membrane of these organelles, and impaired proteolysis capacities^30, 36, 67, 74^. The resulting cholesterol depletion in other intracellular compartments, the potential disorganization in intracellular membranes, as well as the disassembling of essential signaling pathways might globally weaken cell fitness^40, 41, 76, 77^. The loss of NPC1 from the lysosome pool upon MALT1 blockade might notably impact the mTORC1 pathway, again paralleling NPC experimental models^29, 30, 36^. The group of R. Zoncu elegantly demonstrates that NPC1-null cells displayed a hyperactive, detrimental mTORC1 signal^30, 36^. Conversely, we previously described that interrupting MALT1 actions results in the disruption of the mTORC1 pathway^1^. In this line, it has been reported that MALT1 tonic activation may drive the mTORC1 signal, although the underlying mechanisms are not yet fully established^78, 79^. It is tempting to speculate that the resulting mTORC1 hyper- and hypo-activation might vary along with MALT1 status (*i.e.* on-off activity of the enzyme). Nevertheless, the loss of cholesterol transport managed by NPC1 could be rescued by cholesterol supplementation in NPC-defective cell models^30^. Likewise, our data show that the addition of exogenous cholesterol rescues both cell viability and the degradative capacities of lysosomes in MALT-on cells. Whether MALT1 operates upstream of the lysosomal cholesterol-sensing machinery remains to be explored^29–31, 36^. Moreover, the fact that tumor cells relying on MALT1 activity for their survival^53, 68, 69, 79, 80^ exhibits selective sensitivity to NPC1 inhibition raises the possibility that the paracaspase might coordinate the overall organization of the lysosomal compartment toward cholesterol handling.

The Signal Transduction and Activation of RNA (STAR) protein QKI has been shown to regulate GSC fate via the stabilization of miRNA targeting the TGFβ pathway and via the regulation of the endo-lysosomal compartment^1, 2, 81^. Notably, active MALT1 was reported to sequester QKI^1^. Interestingly, two consecutive studies demonstrated the role of QKI in cholesterol homeostasis^82, 83^. The authors provided compelling evidence of the direct binding of QKI to the SREBP2 transcription factor, and further bridging the transcriptional machinery to cholesterol biosynthesis genes^82, 83^. Although our data places SREBP2 activation as a compensatory mechanism, it is alternatively possible that MALT1 blockade, which results in the unleashing of QKI activity^1^, might regulate SREBP2 activation, further reinforcing the concept that MALT1 can orchestrate the cholesterol regulation pathway.

Taken together, our data support the idea that the fitness of MALT1-active cells depends on an effective cholesterol distribution within the cells. These cells are ultimately vulnerable to failure in the cholesterol dispatch, as blocking the NPC1 transporter and/or increasing the number of lysosomes as storage sites prove to be lethal.

## ACKNOWLEDGMENTS

We are grateful to past and present SOAP team members (Nantes Université, INSERM, CNRS, France), especially Agniezska Barbach and Lucas Ottero. We also thank Laetitia Durand and François Paris (Nantes Université, INSERM, CNRS, France). We also thank Cédric Broussard, Johanna Bruce, Philippe Chafey, and François Guillonneau, from Plateforme Proteom’IC 3P5, Université de Paris, Institut Cochin, Paris, France for performing sample preparation, acquisition, and analysis. We would like to acknowledge the core-facilities from UMS Biocore, Nantes, France (MicroPICell ANR-10-INBS-04 and Cytocell).

## AUTHOR CONTRIBUTION

CM, conception and design, acquisition of data, analysis and interpretation of data, draft the article. KT, GAG, MK, LM, KAJ, acquisition of data, analysis and interpretation of data. NB, conception and design, analysis and interpretation of data, draft the article. JG, conception and design, analysis and interpretation of data, write the article. All authors approved the manuscript.

## METHODS

### Cell Culture, Plasmid Transfection, and Lentiviral Transduction

GB patient-derived stem-like cells (GSCs) were obtained as previously described^85^. GSC#1 (mesenchymal, 68-year-old male), GSC#4 (mesenchymal, 76-year-old female), GSC#6 (mesenchymal, 68-year-old male), and GSC#9 (classical, 68-year-old female) were cultured in sphere-forming conditions in NS34 medium (DMEM-F12, Glutamax, and antibiotics, further supplemented with N2 (17502-048), G5 (17503-012), and B27 (17504-044), all from Life Technologies). HEK-293T human embryonic kidney cells, Jurkat E6.1 T lymphocyte cells, BJAB Burkitt lymphoma cells, and SVG-p12 human astrocytes were purchased from the ATCC, OCI-Ly3 B-lymphoma cells were from DSMZ, and all were cultured as per the manufacturer’s instructions. Human brain endothelial cells (hCMEC/D3) were a gift from PO Couraud (Institut Cochin, Paris, France) and cultured accordingly^86^.

pLJC5-Tmem192-3xHA, pLJC5-Tmem192-2xFLAG, pSynSRE-T-Luc, pSynSRE-Mut-T-Luc, and pRL-TK-int-renilla were purchased from Addgene. SRE-T-Luc, SRE-Mut-T-Luc, and renilla plasmid transfection was performed using the GeneJuice transfection reagent following the manufacturer’s instructions (Sigma, 70967). For stable expression of TMEM192-3xHA and TMEM192-2xFLAG, lentiviral particles were produced in HEK-293T cells transfected with pPAX2 and pVSVg, as previously described^87^.

### siRNA Transfection

RNA duplexes targeting the respective human genes were as follows: CAGCAUUCUGGAUUGGCAAAUGGAA (si.2^MALT^^1^), CCUGUGAAAUAGUACUGCACUUACA (si.3^MALT1^), GCGCUCUCAUUUUACCAAATT (si^SREBP2^), ACCAATTGTGATAGCAATATT (si^NPC1^), GGAUGGAGUUAUAAAGGAA (si^NPC2^), and further transfected using RNAiMAX Lipofectamine (Life technologies, 13778150). Stealth non-silencing Low-GC RNA duplexes (si^ctl^, CGACAAUUGUGAGGUCUAAACUAUU, Life Technologies) were used as non-silencing control.

### TCGA and Single Cell RNAseq Database Analysis

The Cancer Genome Atlas (TCGA) was interrogated using the Gliovis Platform (http://gliovis.bioinfo.cnio.es)^58^. RNAseq database was used to investigate data related to SREBP2 and SREBP1 (155 patients) (RNA expression, probability of survival). Agilent 4502A database was used to investigate data related to NPC1 (488 patients) and NPC2 (416 patients) (RNA expression, probability of survival). All subtypes were included. For single cell RNAseq analysis of GB samples, the UCSC Cell Browser was used (https://cells.ucsc.edu)^88^. The glioblastoma infiltrating vs. tumor core dataset^9^ was inquired for SREBP1 and SREBP2 expression.

### Cell Viability Assay

Cell viability was measured using CellTiter-Glo luminescent cell viability assay following the manufacturer’s protocol (Promega, G9243). Experiments were performed in 96-well plates in 100µL final volume. Briefly, GSCs were seeded at 10000 cells per well in triplicate for 2 days with the indicated drugs and vehicle. Alternatively, GSCs were seeded at 8000 cells per well in triplicate for each condition and further challenged with siRNA transfection. Viability was read after 3 more days. Experiments were harvested by the addition of 100µL of the CellTiter-Glo reagent and luminescence was read using a FLUOstar Optima plate reader (BMG).

### Cholesterol/MβCD Complexes Preparation, Antibodies, and Reagents

Cholesterol (Sigma, C3045) was dissolved to a final concentration of 5mM in a solution of 0.1% Mβ D (Sigma, C4555) prepared in sterile H_2_O. The solution was vigorously vortexed, heated at 37°C for 2h, and stored at 4°C.

The following primary antibodies were used: SREBP2 (R&D Systems MAB7119), LDLR (Proteintech 10785-1-AP), GAPDH (Santa Cruz sc-32233), LAMP2 (Santa Cruz sc-18822), P62 (CST #88588), LC3B (CST #3868), HOIL1 (Santa Cruz sc-393754), HA (Sigma-Aldrich H3663), FLAG (Merck F1804), TUBULIN (Santa Cruz, sc-8035), CTSD (BD Biosciences 610800), EEA1 (BD Biosciences 610456), CALRETICULIN (CST 12238), GM130 (Abcam ab52649), VDAC (CST #4661), NPC1 (Abcam ab134113), NPC2 (ABclonal A5413), LBPA (Sigma MABT837). HRP-conjugated secondary antibodies (anti-mouse Ig 1010-05, mouse IgG1 1070-05, mouse IgG2a 1080-05, mouse IgG2b 1090-05, and rabbit 4050-05) were purchased from Southern Biotech. Secondary antibodies for immunofluorescence (Alexa Fluor 546 goat anti-mouse IgG1 A21123, Alexa Fluor 546 goat anti-rabbit A11035, and Alexa Fluor 488 goat anti-mouse IgG1 A21121) were purchased from Thermo Scientific.

The following chemical compounds were used: mepazine (Chembridge, 5216177), MLT748 (Selleckchem, S8898), U18666A (Selleckchem, S9669), cerivastatin (Sigma, SML00005), LLOMe (Sigma, L7393), clemastine (Selleckchem, S1847), and raptinal (Sigma, SML-1745).

### Cell Lysis and Western-blots

Cells were harvested on ice, washed in cold PBS, pelleted (500g, 3min, 4°C) and lysed in RIPA buffer (25mM Tris-HCl pH 7.4, 150mM NaCl, 0.1% SDS, 0.5% Na-Deoxycholate, 1% NP-40, 1mM EDTA) supplemented with Halt protease inhibitor cocktail (Thermo Scientific, #1861279) for 30min on ice. Lysates were cleared by centrifugation (10000g, 10min, 4°C) to pellet insoluble debris and nuclei. Protein concentrations in supernatants were determined using a micro-BCA assay kit (Interchim, 40840A). An equal amount of proteins (10µg) was resolved by SDS-PAGE and transferred onto nitrocellulose membranes (Amersham, 10600002). Proteins were colored and fixed using Ponceau S solution (Santa Cruz, sc-301558), and nonspecific protein binding sites were saturated with 5% milk in PBS-Tween 0.05% (blocking solution). Primary (1/1000 dilution except LAMP2 at 1/5000, GAPDH at 1/20000) and secondary antibodies (1/5000 dilution) were incubated for 1h in the blocking solution. Revelation was performed using Immobilon western chemiluminescent HRP substrate (Merck, WBKLS0500) and the Fusion imaging system (Vilber Lourmat).

### Immunofluorescent Staining

3.10^5^ cells were seeded onto glass slides and fixed for 12min at room temperature with a solution of 4% paraformaldehyde (PFA, Electron Microscopy Sciences, 15710) diluted in PBS. Cells were permeabilized using a solution of Triton-X100 (0.2%, Sigma, T9284) diluted in PBS, for 5min at room temperature. Blocking solution (4% BSA in PBS, Sigma, A2153) was added for 30min prior to incubation 1h at room temperature with primary antibodies (1/200 dilution in the blocking solution). Secondary antibodies (1/400 dilution in the blocking solution) were applied and samples were further processed for confocal analysis.

For dil-LDL uptake, 3.10^5^ cells were treated as indicated (16h for drug treatments, 3 days for siRNA transfection), followed by incubation with dil-LDL (5µg/mL, Khalen biomedical, NC9839048) for 2h at 37°C. Cells were then seeded onto glass slides and fixed for 12min at room temperature with PFA and further processed for confocal analysis.

### Proximity Ligation Assay (PLA)

PLA was performed using the Duolink *in situ* detection reagents far-red kit (Sigma, DUO92013), PLA-probe anti-mouse PLUS (Sigma, DUO92001), and PLA-probe anti-rabbit MINUS (Sigma, DUO92005) according to manufacturer’s protocol. Briefly, GSCs were treated as indicated (3 days siRNA transfection or 6h drug treatment) and seeded onto glass slides before PFA fixation and Triton-X100 permeabilization. Primary antibodies (anti-HA, 1/1000 and anti-NPC1, 1/200) were incubated at 4°C for 16h in a humid chamber before processing according to the manufacturer’s protocol. Samples were processed for confocal analysis.

### Confocal Analysis

Except when mentioned, nuclei were counterstained with DAPI (1/5000, Life Technologies, 62249) and slides were mounted with prolong gold anti-fade mounting medium before imaging (Life Technologies, P36934). Images were acquired on confocal Nikon A1 Rsi, using a 60x oil-immersion lens (IBISA MicroPICell facility, UMS Biocore, Inserm US16, UAR CNRS 3556, Nantes Université, Nantes, France). All images were analyzed and quantified using the ImageJ software.

### Filipin Staining for Imaging and Flow Cytometry

For imagin, cells were seeded onto glass slides and fixed for 12min at room temperature with PFA. PFA was quenched for 10min at room temperature using a glycine/PBS solution (1.5mg/mL, Eurobio, GEPGLY00-66). The filipin III stock solution (25mg/mL DMSO, Sigma, F9765) was diluted to 0.5mg/mL in the blocking solution (BSA 4% in PBS) and added for 2h at room temperature. Finally, cells were mounted with prolong gold anti-fade mounting medium (Life Technologies). Alternatively, cells were incubated with the filipin III/BSA solution for 30min prior to antibody incubation. Primary antibodies were diluted in the blocking solution and incubated for 1h, followed by 30min with secondary antibodies also diluted in the blocking solution. No DAPI counterstaining was performed because the excitation wavelength is the same as filipin III. Slides were quickly imaged on a Zeiss AXIO Observer.Z1.

For flow cytometry analysis, GSCs were similarly processed in 96-V-well plates using a filipin III concentration of 0.125mg/mL diluted in PBS. Fluorescence intensity was measured using the UV 379/28 laser (BD FACSymphony A5, Cytocell facility, UMS Biocore, Inserm US16, UAR CNRS 3556, Nantes Université, Nantes, France). All data were analyzed on FlowJo.

### qPCR

RNA was extracted from 1.10^6^ GSCs using the NucleoSpin RNA Plus purification kit (Macherey-nagel, 740955). Equal amounts of RNA were reverse-transcribed using the Maxima First Strand cDNA Synthesis kit (Thermo Scientific, K1642), and 30ng of the resulting cDNA was amplified by qPCR using PerfeCTa SYBR Green SuperMix Low ROX (VWR, 101414-130). Data were analyzed using the 2-ΔΔCt methods and normalized by the housekeeping genes ACTB and HPRT1.

The following primers targeting the respective human genes were used: ACTB forward 5’-GGACTTCGAGCAAGAGATGG-3’, ACTB reverse 5’- AGCACTGTGTTGGCGTACAG-3’, HPRT1 forward 5’-TGACACTG GCAAAACAATGCA-3’, HPRT1 reverse 5’-GGTCCTTTTCACCAGCAAGCT-3’, HMGCS1 forward 5’-GATGTGGGAATTGTTGCCCTT-3’, HMGCS1 reverse 5’- ATTGTCTCTGTTCCAACTTCCAG-3’, HMGCR forward 5’- GTCATTCCAGCCAAGGTTGT-3’, HMGCR reverse 5’- GGGACCACTTGCTTCCATTA-3’, MVK forward 5’-CATGGCAAGGTAGCACTGG-3’, MVK reverse 5’-GATACCAATGTTGGGTAAGCTGA-3’, FDFT1 forward 5’- ACTTCCCAACGATCTCCCTTG-3’, FDFT1 reverse 5’- CCCATTCTCCGGCAAATGTC-3’, SQLE forward 5’- GATGATGCAGCTATTTTCGAGGC-3’, SQLE Reverse 5’- CCTGAGCAAGGATATTCACGACA-3’, CYP51A1 forward 5’- ATAACCCAGCATCAGGGGAAA-3’, CYP51A1 reverse 5’- CACAGTGGGAAAGTATCCATCAA-3’, FAXDC2 forward 5’- GGCTGCTGACTACATTTGAAGG-3’, FAXDC2 reverse 5’- TCAACCACCAATAGAAGCCCA-3’, NSDHL forward 5’- CAAGTCGCACGGACTCATTTG-3’, NSDHL reverse 5’- ACTGTGCATCTCTTGGCCTG-3’, HSD17B7 forward 5’- ATCTGGACATCATCTCGCAGT-3’, HSD17B7 reverse 5’- AAGAGCTGTAGGGTTCCTTGC-3’, DHCR7 forward 5’- GCTGCAAAATCGCAACCCAA-3’, DHCR7 reverse 5’- GCTCGCCAGTGAAAACCAGT-3’, LDLR forward 5’-AGTTGGCTGCGTTAATGTGAC-3’, LDLR reverse 5’- TGATGGGTTCATCTGACCAGT-3’, ABCA1 forward 5’- GGTGATGTTTCTGACCAATGTGA-3’, ABCA1 reverse 5’- TGTCCTCATACCAGTTGAGAGAC-3’, DGAT1 forward 5’- CCTACCGCGATCTCTACTACTT-3’, DGAT1 reverse 5’- GGGTGAAGAACAGCATCTCAA-3’, NPC1 forward 5’-GTCCAGCGCAGGTGTTTTC- 3’, NPC1 reverse 5’-GCCGAACATCACAACAGAGAC-3’, NPC2 forward 5’- CAAAGGACAGTCTTACAGCGT-3’, NPC2 reverse 5’- GGATAGGGCAGTTAATTCCACTC-3’.

### Luciferase SREBP2 Reporter Assay

2.10^6^ cells were transfected with 2µg of either pSynSRE-T-Luc or pSynSRE-Mut-T-Luc in combination with 0.1µg pRL-TK-Renilla using the GeneJuice transfection reagent. After 24h, cells were seeded in a 96-well plate in triplicate per condition and cultured for a further 16h in the presence of the indicated drugs. At the end of the experiment, cells were pelleted and lysed in 30µL of lysis buffer. 20µL of the lysate was revealed using the Dual-Glo luciferase assay system following the manufacturer’s protocol (Promega, E1910). Luminescence was measured using a FLUOstar Optima plate reader (BMG).

### Cholesterol Dosage

Cellular cholesterol was measured using the Cholesterol/Cholesterol Ester-Glo Assay Kit (Promega, J3190) following manufacturer’s instructions. Briefly, 1.10^5^ GSCs were lysed for 30min at 37°C. A volume of 25µL of the lysate was used for cholesterol quantification. Cholesterol level was determined by reading luminescence after 1h incubation with cholesterol reductase and cholesterol esterase reagents at room temperature. Cholesterol concentration was extrapolated from a standard curve prepared for each experiment and normalized to protein concentration.

### Lysosome Immunoprecipitation (LysoIP)

15.10^6^ GSCs expressing TMEM192-3xHA were used per condition. TMEM192-2xFLAG expressing cells were used as a control. Each step was conducted at 4°C. After the indicated treatments, cells were washed in cold PBS and centrifuged (1000g, 2min). Pelleted cells were resuspended in 500µL cold PBS + anti-proteases, and 100µL was saved for whole cell lysate control. The remaining 400µL was mechanically lysed with 10 strokes of a 29-gauge syringe and centrifuged (1000g, 2min). Supernatants containing organelles were incubated with 75µL of Pierce anti-HA magnetic beads (Thermo Scientific, 88837) for 15min on a rotator wheel. Beads were then washed 3 times with cold PBS + anti-proteases. For further analysis, dried beads were eluted twice with 50µL of elution buffer (50mM Tris/HCl pH 8.5 with 2% SDS). Eluates were denaturated (95°C, 5min) and further processed for western-blot and proteomic analysis. Alternatively, samples were stored at -80°C.

### Label-Free Quantification (LFQ) Proteomic Processing and Analysis of Whole Cell Lysates

For sample preparation, pelleted cells were solubilized in lysis buffer (2% SDS, 200mM TEAB, pH 8.5) and heated for 5min at 95°C. The protein concentration of the supernatants was estimated by BCA assay. Proteins were then reduced and alkylated with 10mM TCEP and 50mM chloroacetamide. Bottom-up experiments’ tryptic peptides were obtained by S-Trap Micro Spin Column according to the manufacturer’s protocol (Protifi, NY, USA). Briefly, 30μg of proteins were digested with 1µg Trypsin sequencing grade (Promega) for 14h at 37°C. The S-Trap Micro Spin Column was used according to the manufacturer’s protocol. After speed-vacuum drying, eluted peptides were solubilized in 2% trifluoroacetic acid (TFA) and fractionated by strong cationic exchange (SCX) Stage-Tips, as previously described^89^.

Liquid Chromatography-coupled Mass spectrometry analysis (LC-MS) analyses were performed on a Dionex U3000 RSLC nano-LC-system (Thermo Fisher scientific, Les Ulis, France) coupled to a TIMS-TOF Pro mass spectrometer (Bruker Daltonik GmbH, Bremen, Germany). After drying, peptides from SCX Stage-Tip, the 5 fractions were solubilized in 10μL of 0.1% TFA containing 10% acetonitrile. One μ was loaded, concentrated, and washed for 3min on a C_18_ reverse phase precolumn (3μm particle size, 100 Å pore size, 75 μm inner diameter, 2 cm length, from Thermo Fisher Scientific). Peptides were separated on an Aurora C_18_ reverse phase resin (1.6μm particle size, 100Å pore size, 75μm inner diameter, 25cm length mounted to the Captive nanoSpray Ionisation module, IonOpticks, Middle Camberwell Australia) with a 120-minutes overall run time with a gradient ranging from 99% of solvent A containing 0.1% formic acid in milliQ-grade H_2_O to 40% of solvent B containing 80% acetonitrile, 0.085% formic acid in mQH_2_O. The mass spectrometer acquired data throughout the elution process and operated in DDA PASEF mode with a 1.1 second/cycle, with Timed Ion Mobility Spectrometry (TIMS) mode enabled and a data-dependent scheme with full MS scans in Parallel Accumulation and Serial Fragmentation (PASEF) mode. This enabled a recurrent loop analysis of a maximum of the 120 most intense nLC-eluting peptides which were CID-fragmented between each full scan every 1.1sec. Ion accumulation and ramp time in the dual TIMS analyzer were set to 166 msec each and the ion mobility range was set from 1/K0 = 0.6 Vs cm^-2^ to 1.6 Vs cm^-2^. Precursor ions for MS/MS analysis were isolated in positive mode with the PASEF mode set to « on » in the 100-1.700 m/z range by synchronizing quadrupole switching events with the precursor elution profile from the TIMS device. Singly charged precursor ions were excluded from the TIMS stage by tuning the TIMS using the otof control software (Bruker Daltonik GmbH). Precursors for MS/MS were picked from an intensity threshold of 1000 arbitrary units (a.u.) and resequenced until reaching a ‘target value’ of 20.000 a.u. taking into account a dynamic exclusion of 0.40 s elution gap.

Regarding protein quantification and comparison, mass spectrometry data were analyzed using Maxquant version 1.6.6.0^90^. The database used was a *Human* sequence from the Uniprot databases (release March 2020). The enzyme specificity was that for trypsin. The cleavage specificity was trypsin’s with maximum 2 missed cleavages. Carbamidomethylation of cysteines was set as constant modification, whereas acetylation of the protein N terminus and oxidation of methionines were set as variable modifications. The false discovery rate was kept below 1% on both peptides and proteins. LFQ was performed using both unique and razor peptides. At least two such peptides were required for LFQ. The “match between runs” (MBR) option was allowed with a match time window of 0.7min and an alignment time window of 20min. For differential analysis, LFQ results from MaxQuant were imported into Perseus software version 1.6.14.0 (Max-Planck Institute of Biochemistry). Reverse and contaminant proteins were excluded from the bioinformatic analysis. LFQ data were transformed into log_2_. A t-test (p-value<0.05) was carried out on proteins, with 3 valid values in at least one group. Moreover, PCA (principal component analysis) was performed with imputation.

### Label-Free Quantification (LFQ) Proteomic Processing and Analysis of LysoIP

For sample preparation, IP samples were solubilized in lysis buffer (2% SDS, 200mM Tris-HCl, pH 8.5, 10mM TCEP, 50mM chloroacetamide). Bottom-up experiments’ tryptic peptides were obtained by S-Trap Micro Spin Column according to the manufacturer’s protocol (Protifi, NY, USA). Proteins were digested with 1µg Trypsin sequencing grade (Promega) for 14h at 37°C. The S-Trap Micro Spin Column was used according to the manufacturer’s protocol. After speed-vacuum drying, eluted peptides were solubilized in 10µL of 10% acetonitrile and 0.1% trifluoroacetic acid. LC-MS analysis was done with similar parameters than the proteome-wide LFQ analysis (see previous section), except the peptide separation run time was 60min, ion accumulation and ramp time was 100ms, and precursors picking for MS/MS: 2500 arbitrary units (a.u.). The parameters for protein quantification and comparison were the same as described in the previous section.

### RNAseq Analysis

5.10^6^ GSC#9 were treated for 4h with vehicle (DMSO) and MPZ (20µM) and snap-frozen on dry ice. Samples and data were processed at Active Motif (Carlsbad, California, USA) as previously described^1^.

### Proteomic and RNAseq Enrichment Analysis

Enrichment analysis of the proteomic and RNAseq experiments were performed using g:Profiler^91^ (version e107_eg54_p17_bf42210) applying a significance threshold of 0.05. The Panther classification system^84^ was also used to identify the main GO:function enriched in the proteomic analysis.

### Statistical Analysis and Quantification

Densitometry and imaging quantifications were performed using the ImageJ software. All graphs were mounted and statistically tested using Prism 8 (GraphPad). Unless otherwise specified, error bars on graphs are shown as mean ± s.d. of at least three independent experiments. A p-value of <0.05 was considered significant. The RNAseq experiment was performed on three independent biological replicates. All proteomics experiments were performed on four independent biological replicates. Viability assays were performed on three independent experiments, each in triplicate.

## Figure Legends to Supplemental Tables

**Table S1. Proteome-wide Label-Free Quantification in Control Versus Mepazine-treated GSC. Supplemental Table to Figures 1, 4, and S4**.

Statistical and fold-change analysis, as well as Principal Component Analysis (PCA) and heatmaps were done with Perseus software (1.6.14.0) to compare DMSO and mepazine (MPZ)-treated groups. Each group was performed in quadruplicates.

**Table S1. List of Upregulated GO Term Signatures in the WCL Proteomic Analysis of MALT1-inhibited GSCs. Supplemental Table to Figure 1**.

Differentially up-regulated proteins from DMSO versus MPZ-treated GSC#9 were analyzed with the Pantherdb^84^ pathway browser. Four main GO term signatures were identified as follows: 1. actin (1.a: actin filament network formation, 1.b: actin filament bundle assembly, 1.c: actin filament bundle organization, 1.d: positive regulation of actin filament polymerization), 2. spindle (2.a mitotic spindle organization, 2.b: mitotic spindle organization, 2.c: microtubule cytoskeleton organization involved in mitosis, 2.d: spindle assembly, 2.e: spindle organization), 3. lipid (3.a fatty acid oxidation, 3.b lipid oxidation), and, 4. RNA (4.a transcription elongation from RNA polymerase II promoter).

**Table S3. Immunopurified Lysosomes Proteome Label-Free Quantification in Control Versus Mepazine-treated GSC. Supplemental Table to Figures 4 and S4.**

Statistical and fold change analysis, as well as Principal Component Analysis (PCA) and heatmaps were done with Perseus software (1.6.14.0) to compare the following groups: HA_DMSO vs Flag_DMSO, HA_MPZ vs Flag_MPZ, Flag_MPZ vs Flag_DMSO and HA_MPZ vs HA_DMSO. Statistical and fold-change analysis were done with LFQ values (without imputation) and paired t-test p-value. Each group was performed in quadruplicates.

## Figure Legends to Supplemental Figures

**Figure S1. SREBP2 is Preferentially Activated in Response to MALT1 Inhibition. Supplemental Information to Figure 1**

(A-B) (Top) Violin plot of SREBP2 (A) and SREBP1 (B) mRNA expression in non-tumor and GB samples from the TCGA RNAseq dataset. Each dot represents one clinical sample. (Bottom) Kaplan-Meier curve of the probability of survival for >77 GB patients with low (green) or high (red) SREBP2 RNA level based on the TCGA RNAseq dataset.

(C) (Left) 2D-tSNE representation of cell clusters identified in 4 different GB samples^9^ using the UCSC portal^88^. One dot refers to one cell. (Right) SREBP2 and SREBP1 expression pattern across the 2D-tSNE space. The relative expression level of the respective targets is indicated.

(D) (Left) Densitometric analysis of the level of cleaved fragment of SREBP2 normalized to its full-length form, from representative western-blots shown in panel 1I. The value was then normalized to GAPDH level. GSC#4, GSC#6, and GSC#9 were analyzed. (Right) Densitometric analysis of LDLR level normalized to GAPDH, from representative western-blots shown in panel 1I. GSC#4, GSC#6, and GSC#9 were analyzed. Data are presented as the mean ± s.d. of 4 independent experiments.

(E) qRT-PCR of DGAT1 in GSC#9 treated for 6h with vehicle (DMSO), mepazine (MPZ, 20µM), and MLT748 (5µM). Alternatively, cells received non-silencing RNA duplexes (si^ctl^) and 2 independent RNA duplexes targeting MALT1 (si.2^MALT1^ and si.3^MALT1^) for 3 days. Data were normalized to 2 housekeeping genes (HPRT1, ACTB) and are presented as the mean ± s.d. of 3 independent biological replicates.

All panels are representative of at least n=3, unless otherwise specified. t-test and ANOVA, *p<0.05, **p<0.01, ***p<0.001.

**Figure S2. SREBP2 Blockade Exacerbates MALT1-inhibition Induced Phenotypes. Supplemental Information to Figure 2**

(A) Western-blot analysis of the levels of SREBP2 and LDLR in GSC#9 transfected for 2 days with non-silencing (si^ctl^) and SREBP2 targeting (si^SREBP2^) RNA duplexes, and further treated for 24h with vehicle (DMSO) and mepazine (MPZ, 20µM). GAPDH served as a loading control.

(B) Cell viability was measured using CellTiter-Glo assay in GSC#9 transfected as described in (A), and further treated for 48h with DMSO and MPZ (20µM). Data are presented as the mean ± s.d. of at least 3 independent biological replicates.

(C) Cell viability was measured using CellTiter-Glo assay in GSC#9 that were pre-treated 1h with DMSO and cerivastatin (10nM), and then further challenged for 48h with DMSO and MPZ (20µM). Data are presented as the mean ± s.d. of at least 3 independent biological replicates.

(D) Western-blot analysis of the levels of SREBP2, P62, and LC3B in GSC#9 treated as described in (A). GAPDH served as a loading control. Unlipidated (LC3B-I) and lipidated (LC3B-II) LC3B forms are indicated with red and green arrowheads, respectively.

(E) (Left) Confocal analysis of P62 staining (red) in GSC#9 treated as described in (A). Nuclei are shown in blue (DAPI). Scale bar: 10µm. (Right) Violin representation of the number of P62 punctae per cell, n>69.

(F) qRT-PCR of ABCA1 in GSC#9 treated for 4h with DMSO and MALT1 inhibitors (MPZ, 20µM, and MLT748, 5µM). Alternatively, cells received si^ctl^, si.2^MALT1^, and si.3^MALT1^, for 3 days. Data were normalized to 2 housekeeping genes (HPRT1, ACTB) and are presented as the mean ± s.d. of 3 independent biological replicates. All panels are representative of at least n=3, unless otherwise specified. t-test and ANOVA, *p<0.05, **p<0.01, ***p<0.001.

**Figure S3. MALT1 Activity is not Dependent on Cholesterol Concentration. Supplemental Information to Figure 3**

(A) Western-blot analysis of the level of LDLR in GSC#9 pre-treated for 1h with DMSO and MALT1 inhibitors (MPZ, 20µM, and MLT748, 5µM), and further challenged for 24h with vehicle (H_2_O) or with MβCD-complexed cholesterol (chol/MβCD, 250µM). GAPDH served as a loading control.

(B) Western-blot analysis of the level of HOIL1 in GSC#9 treated as in (A). GAPDH served as a loading control. FL (full length) and cleaved HOIL1 forms are indicated with red and green arrowheads, respectively.

(C) Confocal analysis of TMEM192-3xHA (HA) staining (red) in HA-lyso GSC#9 treated as described in (A). Nuclei are shown in blue (DAPI). Scale bar: 10µm.

All panels are representative of at least n=3, unless otherwise specified. t-test and ANOVA, *p<0.05, **p<0.01, ***p<0.001.

**Figure S4. Lysosomes from MALT1-Inhibited GSCs Accumulates Autophagic Cargo. Supplemental Information to Figure 4**

(A) Graph represents the number of proteins identified in the whole cell lysate (WCL) and the immunopurified lysosome fractions (grey bar graph), as depicted in panel 4A. The percentage of GO:CC lysosomes proteins is indicated in green. Donut graph represents the percentage of membranous (blue) and luminal (green) lysosomal proteins identified in the LysoIP proteomic analysis. The percentage of other type of proteins is shown in grey. For each graph, the number of proteins is reported.

(B) GO:BP enrichment analysis of proteins differentially expressed with fold change >1.5, either positively (red bar graphs) and negatively (blue bar graphs), in immunopurified lysosomes from vehicle (DMSO) and mepazine (MPZ, 20µM) treated HA-lyso GSC#9.

(C) Volcano plot analysis of differentially expressed proteins from (B). GO:BP autophagy associated proteins are identified (yellow dots).

(D) qRT-PCR of NPC1 and NPC2 in GSC#9 treated for 6h with vehicle (DMSO) and MALT1 inhibitors (MPZ, 20µM, and MLT748, 5µM). Alternatively, cells received non-silencing (si^ctl^) and MALT1 targeting (si.2^MALT1^ and si.3^MALT1^) RNA duplexes for 3 days. Data were normalized to 2 housekeeping genes (HPRT1, ACTB) and are presented as the mean ± s.d. of 3 independent biological replicates.

All panels are representative of at least n=3, unless otherwise specified. t-test and ANOVA, *p<0.05, **p<0.01, ***p<0.001.

**Figure S5. NPC1 Inhibition Selectively Abrogates GSC Survival. Supplemental Information to Figure 5**

(A) Kaplan-Meier curve of the probability of survival for >243 GB patients with low (green) and high (red) *NPC2* RNA level based on the TCGA Agilent-4502A dataset.

(B) Western-blot analysis of the levels of NPC2, LAMP2, and P62 in GSC#9, transfected with non-silencing (si^ctl^) and NPC2 targeting (si^NPC2^) RNA duplexes for 3 days. GAPDH served as a loading control.

(C) Cell viability was measured using CellTiter-Glo assay in each cell line treated for 48h with DMSO and U18666A at the indicated doses. Data are presented as the mean ± s.d. of at least 3 independent biological replicates.

(D) Heatmap of the cell viability obtained in (C).

All panels are representative of at least n=3, unless otherwise specified. t-test and ANOVA, *p<0.05, **p<0.01, ***p<0.001.

