## Supplementary figures and images for "The paracaspase MALT1 controls cholesterol homeostasis in glioblastoma stem-like cells through lysosome proteome shaping"

### Supplemental FIgure

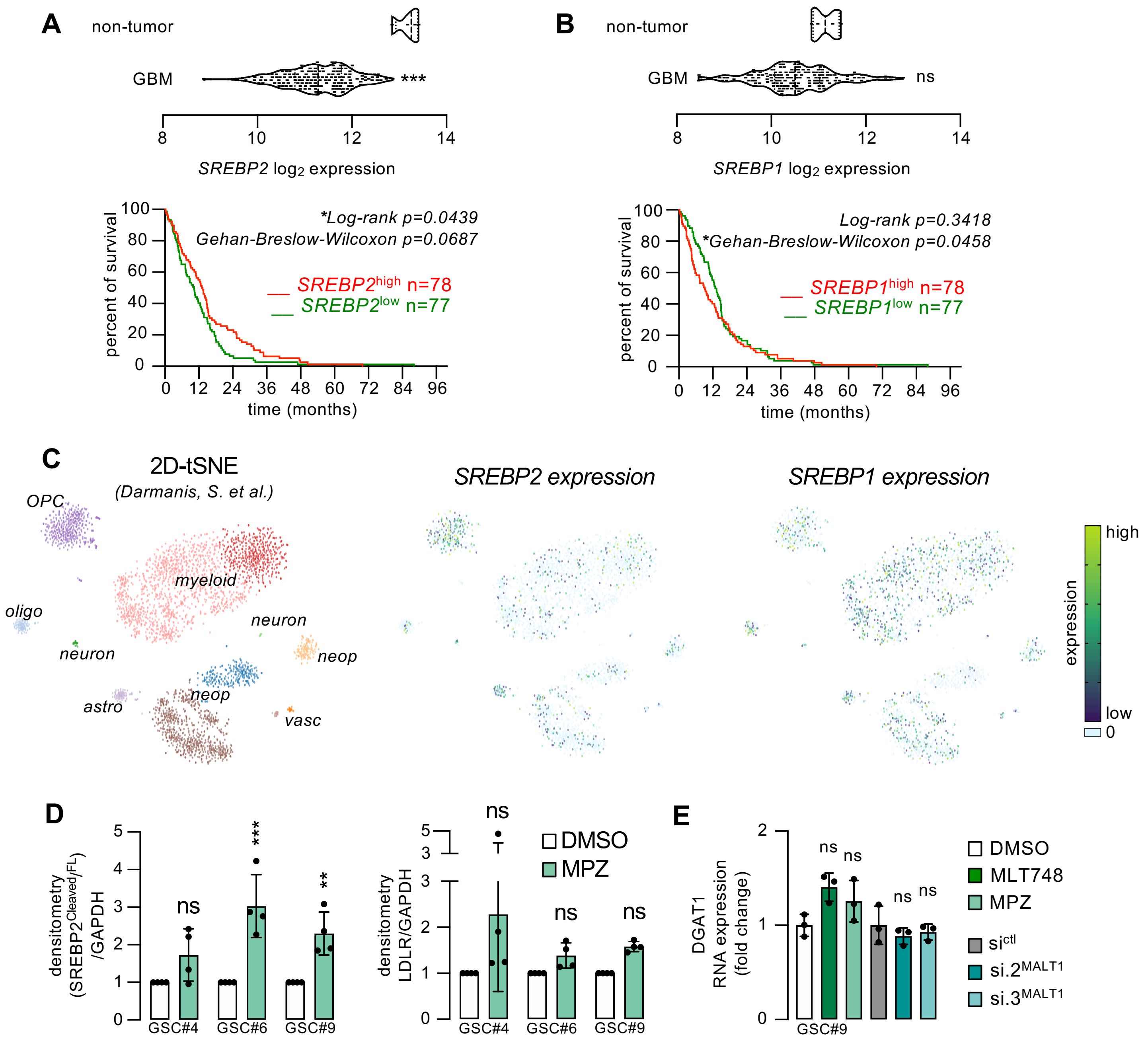

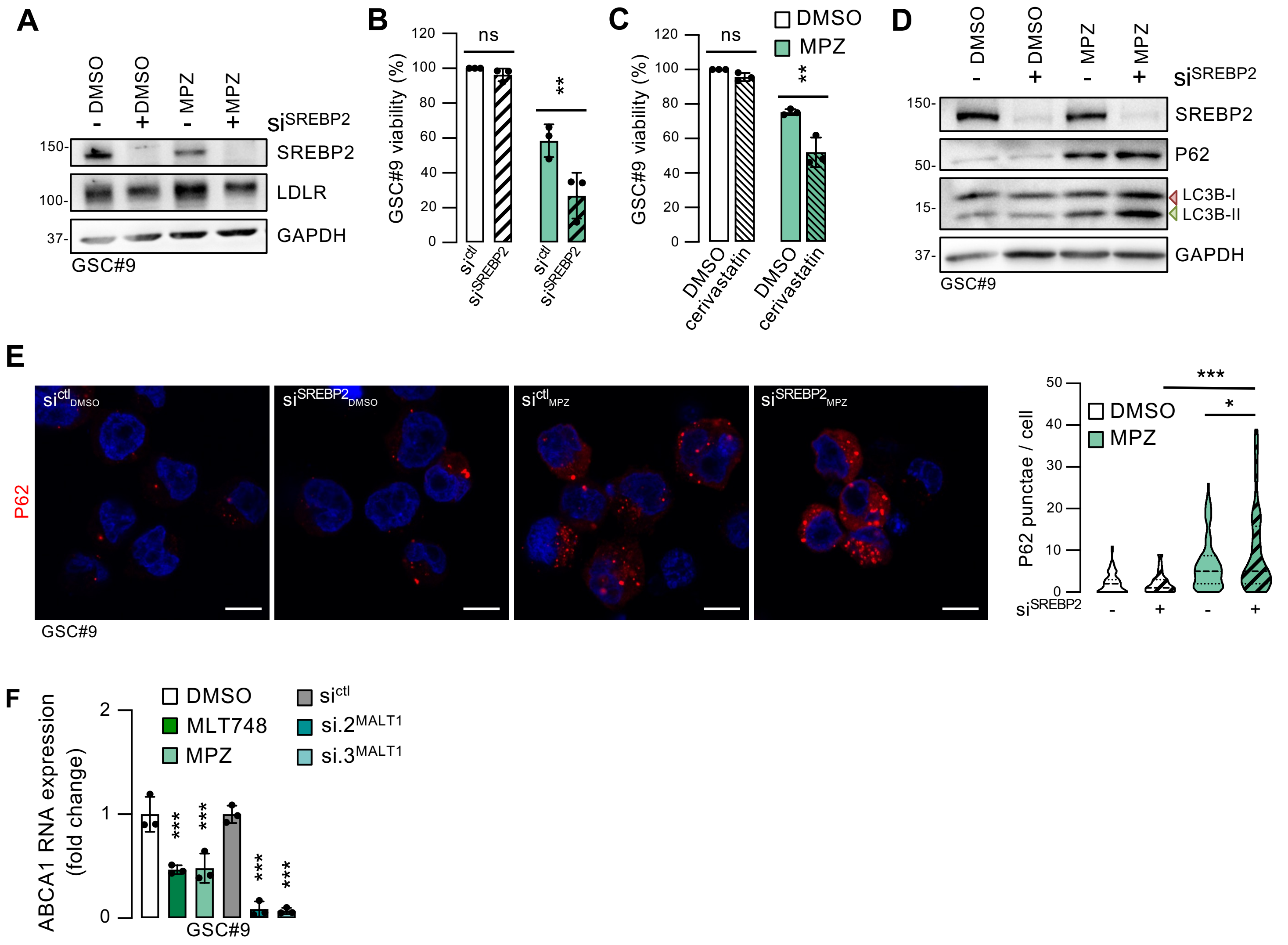

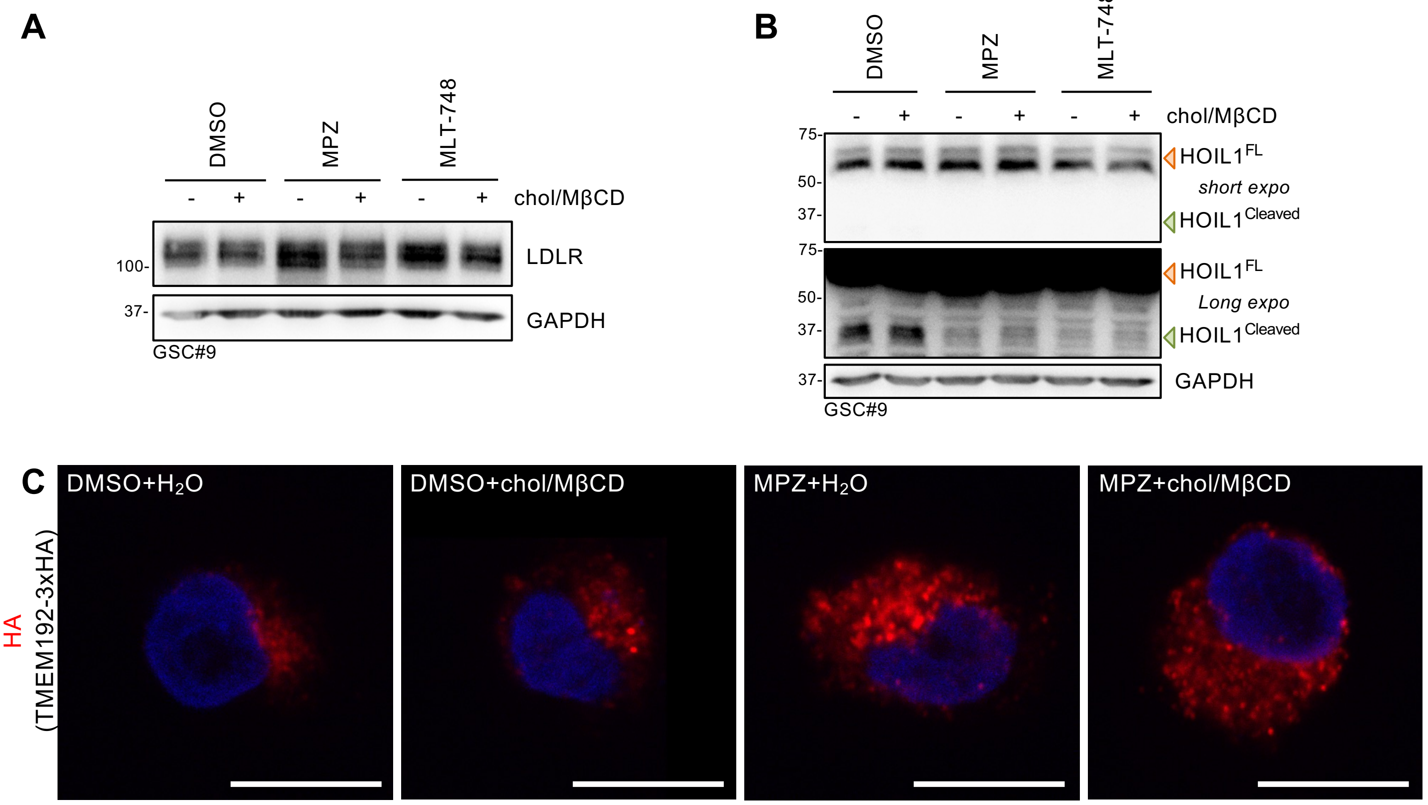

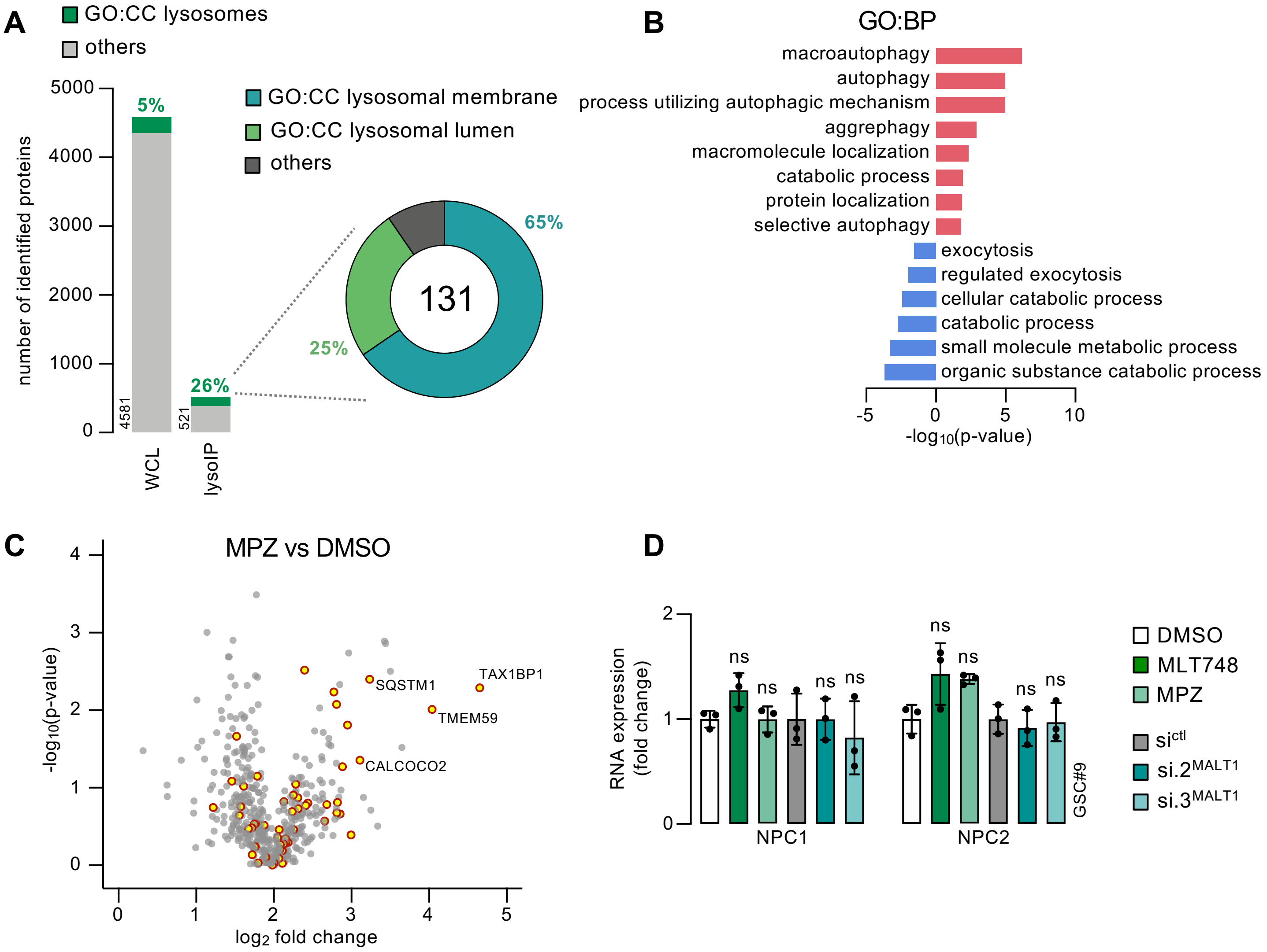

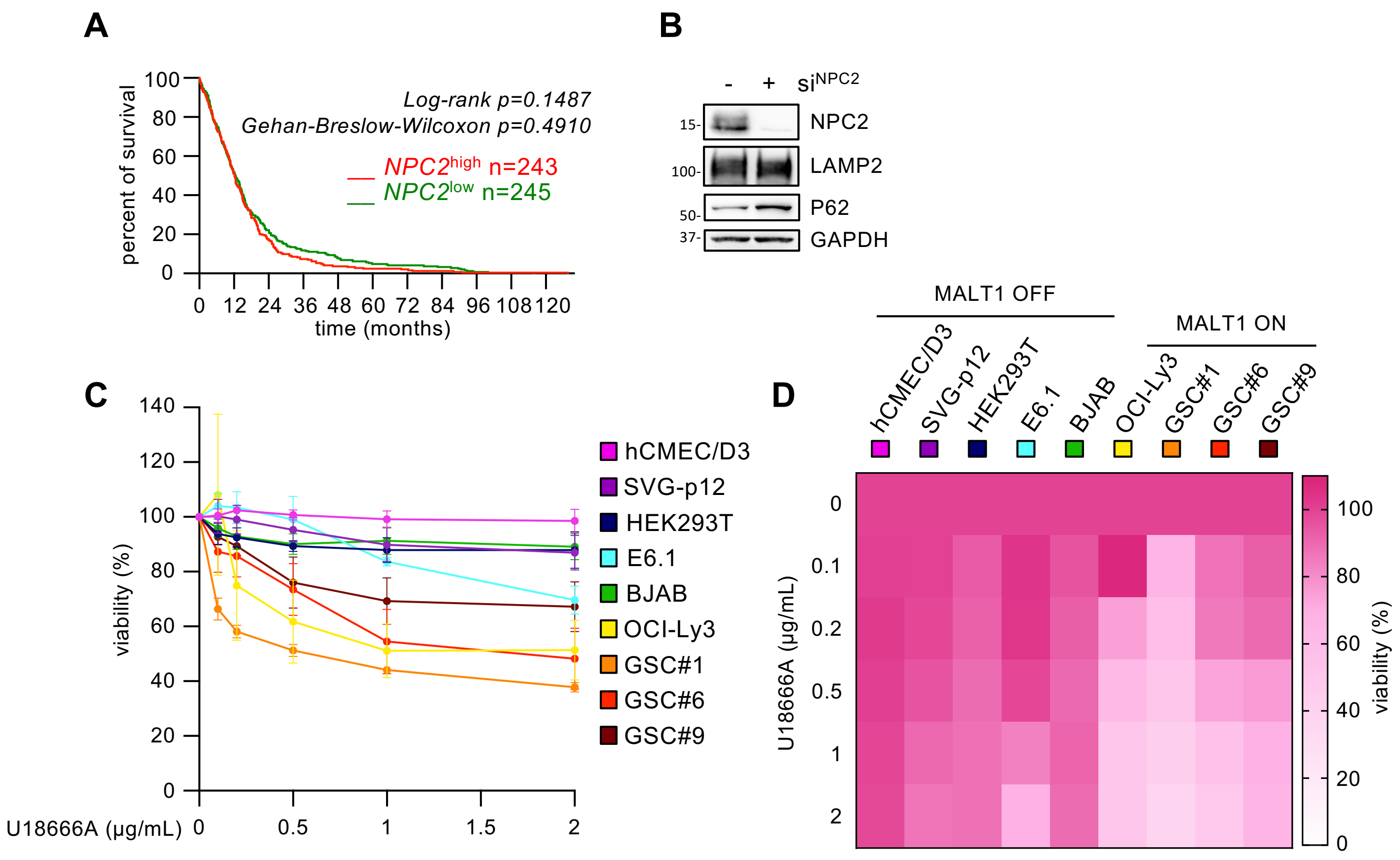
